# Comparative whole-genome analysis of a Thar desert strain *Streptomyces* sp. JB150 provides deep insights into the encoded parvome and adaptations to desert edaphic system

**DOI:** 10.1101/2021.01.20.427384

**Authors:** Dharmesh Harwani, Jyotsna Begani, Jyoti Lakhani

## Abstract

We sequenced the genome of *Streptomyces* sp. JB150, isolated from a unique site of the Thar desert in India. Genome mining of the JB150 genome revealed the presence of many interesting secondary metabolic biosynthetic gene clusters (BGCs). The encoded parvome of JB150 includes non-ribosomal peptides, polyketides including β-lactone, butyrolactone, ectoine, lantipeptides, lasso peptides, melanin, resorcinol, siderophores, terpenoids, thiopeptides, and other types of hybrid compounds. Among them, ~30% BGCs displayed a high degree of novelty. The genome of JB150 was enriched for a large assortment of specialized genes coding for the production of many interesting biomolecules comprising compatible solutes, multiple stress-response regulators, transport proteins, protein secretion systems, signaling molecules, chaperones and storage reserves, etc. The presence of diverse members of CAZymes enzyme families, high numbers of riboflavin, flavin mononucleotide (FMN) and flavin adenine dinucleotide (FAD), trehalose and aromatic compounds synthesis genes, putative orthologues to several of the classical fatty acid synthesis components, prototrophy for many essential amino acids exhibit metabolic versatility of JB150 to inhabit in the extreme desert environment. Besides, the genome of JB150 was observed to specifically encode thiazole-oxazole-modified thiazolemicrocin (TOMM) and ectoine. The comparison of the complete genomes of *Streptomyces* sp. JB150 and seven other actinomycete strains belonging to different desert ecosystems unveiled the presence of many previously undetected, distinctive, biological, and genomic signatures. We propose that these genetic traits endowed by these strains are essential for their adaptation in the highly underprivileged, extreme ecosystem of the Thar desert to cope with multiple abiotic stressors, oligotrophic nutrient conditions and to produce a huge repertoire of diverse secondary metabolites.

## Introduction

Actinomycetes, the most primitive lineages among prokaryotes, are believed to evolve about 2.7 billion years ago [1,2]. It contains several genera including *Streptomyces* which harbor diverse secondary metabolic pathways and produce around two-thirds of all known antibiotics [3,4]. *Streptomyces* have linear chromosomes and their genome size ranges from 6.2 Mb in *Streptomyces cattleya* to 12.7 Mb in *Streptomyces rapamycinicus* [5,6]. Although the species of this industrially important genus are widely distributed, yet the rate of the discovery of novel natural products produced by them has been drastically reduced. Furthermore, an alarmingly high rate of the development of multidrug-resistance in microbial pathogens necessitates the search for yet undiscovered and metabolically potent actinomycete species from specialized niches such as deserts for potential new secondary metabolites [7–9]. In the present work, we report the draft genome sequence of *Streptomyces* sp. JB150. The strain JB150 was isolated from the Bikaner region of the Thar desert which is known as the tropical desert of Asia that extends to India through Rajasthan and Gujarat. For comparison, seven other complete genomes of desert-actinomycetes: *Actinoalloteichus spitiensis* RMV-1378 (cold desert, Indian Himalaya) [10], *Geodermatophilus africanus* DSM 45422 (Sahara desert, Ourba) [11], *Jiangella gansuensis* DSM 44835 (Gansu desert, China) [12], *Lechevalieria deserti* DSM 45480 (Atacama desert, Chile) [13], *Modestobacter excelsi* 1G6 DSM 107535 (Atacama desert, Chile) [14], *Streptomyces* sp. DH-12 (Thar desert, India) [15], and *Streptomyces* sp. Wb2n-11 (Sinai desert, Egypt) [16] and were also included in the present study. These taxonomical relatives of JB150 belonging to the different desert ecosystems were purposefully selected to investigate the genetic assortment and similarities that confer upon them the ability to sustain and thrive under the harsh conditions of deserts and produce a huge repertoire of diverse secondary metabolites.

The genome investigation of JB150 unveils the presence of a large assortment of specialized genes coding for the production of many interesting biomolecules comprising secondary metabolites, stress response regulators, transport proteins, protein secretion systems, signaling molecules, storage reserves, etc. The thermal adaptation in JB150 may partially be attributed to the presence of relatively high GC content compared to other analyzed actinomycetes strains. The genome analysis revealed the presence of 24 biosynthetic gene clusters (BGCs) involved in the production of different secondary metabolites such as non-ribosomal peptides, polyketides including β-lactone, butyrolactone, ectoine, lantipeptide, lasso peptide, melanin, resorcinol, siderophore, terpenoids, thiopeptide, and several types of hybrid compounds. Among these, 7 BGCs, exhibit no or very less relative similarity to already known clusters, possibly encode for the novel metabolic compounds. To regulate osmotolerance, the strain JB150 possesses the genetic machinery to produce compatible solutes such as ectoine and proline. To resist the detrimental effects of the high osmolarity in the desert environment, JB150 bears two glycine betaine uptake systems as well. The proteins for well-defined defense mechanisms against oxidative and nitrosative stress and a stress-responsive alternative sigma factor were also prominent in JB150. Furthermore, to sustain under oxygen limitation or oxidative stress, the strain contains hemoglobin-like favohemoproteins. Several chaperones were also identified in JB150 that may play their roles in the transit of outer membrane proteins through the periplasm. To mediate communication between the intracellular and the extracellular environment JB150 genome contains a total of 591 membrane transporter families most of which belong to ATP-binding cassette (ABC) and major facilitator superfamily (MFS) transporters. The presence of twin-arginine targeting (Tat) family and other transporters in JB150 may also play their roles in acquiring nutrients efficiently in the desert environment. Similarly, JB150 also possesses Na^+^: H^+^ antiporters (NhaA), which could further assist the strain to maintain the pH homeostasis by facilitating removal of excessive Na^+^ to prevent toxicity in the cell. The genome of JB150 also contains the putative components of type V and type VII secretion systems.

Compared to other genomes, it appears that JB150 has the ability to synthesize the majority of amino acids. It comprises the highest number of genes implicated in methionine, threonine, and homoserine biosynthesis. Prototrophy for many essential amino acids and the capacity to grow without needing external sources might be advantageous to the strain in nutrient-poor soil environments. Additionally, the presence of 1,3-diaminopropane (DAP) pathway and a high number of genes for the synthesis of aromatic compounds and proline could also contribute massively in the biosynthesis of antibiotics, defense molecules, siderophores, pigments, signaling compounds, and various secondary metabolites in JB150. Intriguingly, among the analyzed genomes, the highest number of trehalose biosynthesis genes was identified in JB150. The accumulation of trehalose in high amounts might play a significant role in the strain to gain an additional advantage in the high temperature and oligotrophic nutrient conditions. The presence of diverse members of polysaccharide lyases (PL) and other CAZymes enzyme families, major membrane phospholipids such as phosphatidylethanolamine (PE), phosphatidylglycerol (PG), cardiolipin (CL), primary storage lipophilic compounds like polyhydroxyalkanoates (PHA) and triacylglycerols (TAG), complete set of enzymes required for DNA replication, recombination, and DNA repair, unusually high numbers of riboflavin, FMN and FAD genes and putative orthologues to several of the classical FAS II components may further assist JB150 to survive and sustain in the arid desert environment. Besides, the comparison of BGCs involved in the secondary metabolism among the analyzed genomes unveils the presence of a unique gene cluster for thiazole-oxazole-modified thiazolemicrocin (TOMM) biosynthesis in JB150. Moreover, the presence of a high number of genetic determinants to regulate cation transporters, 4-hydroxyproline uptake and utilization, ectoine biosynthesis, oxidative stress, pentose phosphate pathway in JB150 relative to other analyzed genomes might be advantageous to respond multiple stresses in desert soils, characterized by low nutrient and high salinity. Several comparisons among JB150 and other seven selected genomes revealed enrichment of commonly shared genes which may have potential roles in secondary metabolite synthesis, stress-response regulation, membrane transport, nitrogen metabolism, DNA metabolism, and in the synthesis of betaine, glutathione, cofactors, vitamins, prosthetic groups, and pigments. Apart from possessing the classical sets of housekeeping genes, many other biosynthetic gene clusters (BGCs), occupied in the biosynthesis of fatty acids, lipids, isoprenoids as well as clusters of orthologous groups (COGs) of proteins, carbohydrate-active enZYmes (CAZymes) and protein families marked their obvious presence in JB150 and other 7 analyzed genomes. We propose that the genetic traits endowed by these strains are essential for their adaptation in the highly underprivileged desert conditions to cope with multiple abiotic stressors, oligotrophic nutrient conditions and to produce a huge repertoire of diverse secondary metabolites.

## Material and methods

### Selective isolation and culture media

For the isolation of actinomycetes, the soil sample was collected at Bikaner (28.0229°N, 73.3119°E) region falling in the arid Thar desert in the Indian state Rajasthan. The streptomycete JB150 was selectively isolated, using *<u>M</u>*odified *<u>A</u>*ctinomycete *<u>S</u>*elective (MAS)-DH agar medium (composition per liter; 1g starch, 1g sodium caseinate, 0.1g L-asparagine, 4g sodium propionate, 0.5g dipotassium phosphate, 1g potassium nitrate, 0.1g magnesium sulfate, 0.001g ferrous sulfate, 0.02g calcium carbonate, 20g agar). The visualization of the isolate was done using a bright-field microscope (1000x) and a mobile phone mounted Foldscope-microscope (140x) [17]. The pure culture of the isolate was stored at −40°C using 20% (w/v) sterile glycerol until further use.

### Phylogenetic analysis

The genomic DNA of each isolate was extracted using a method of Li *et al.* [18]. The 16S rDNA of the isolate JB50 was amplified using PCR with universal primers 27F 5’AGA GTT TGA TCC TGG CTC AG 3’ and 1525R 5’ AAG GAG GTG ATC CAG CCG CA 3’ that will amplify specifically 16SrDNA region [19–21]. PCR-amplicons were visualized in 2% agarose gel electrophoresis and subsequently revealed with ethidium bromide staining. The amplified 16SrDNA gene product was sequenced using the Sanger Didoxy method. Manual sequence edition, alignment, and contig assembling were performed using the Vector NTI *v 10* software package. Sequencing results were analyzed for chimeras using the DECIPHER *v 1.4.0* program [22]. The 16S rDNA sequence of JB150 was deposited in NCBI under the sequences accession number MT649861. Reference sequences were downloaded from the Genbank Database (http://www.ncbi.nlm.nih.gov/genbank). To assess the taxonomical relatedness between *Streptomyces* sp. JB150 and 7 other complete genomes of *Actinoalloteichus spitiensis*, *Geodermatophilus africanus*, *Jiangella gansuensis, Lechevalieria deserti*, and *Modestobacter excelsi* 1G6, *Streptomyces* sp. DH-12 and *Streptomyces* sp. Wb2n-11, a phylogenetic analysis employing concatenated sequences of *gyrA* (DNA gyrase subunit A), *rpoB* (DNA-directed RNA polymerase subunit beta), and *rsmH* (ribosomal RNA small subunit) was also performed. The sequences were aligned by using the multiple sequence alignment program ClustalW. The Neighbor-Joining algorithm [23] was used with a Kimura distance matrix model in MEGA X. All nodes supported by 1000 bootstrap replications were used to construct a phylogenetic tree [24].

### DNA sequencing and assembly

The genomic DNA was isolated from *Streptomyces* sp. JB150 as described above. The quality and quantity of the genomic DNA were assessed on Agarose gel and Nano-drop One (Thermo Fisher Scientific). The paired-end (PE) 2×150bp NextSeq500 shotgun library was prepared using Illumina TruSeq *v3.0* genomic DNA library preparation kit. The library was loaded on Illumina Next Seq 500 system for cluster generation and sequencing. PE sequencing allowed the template fragments to be sequenced in both the forward and reverse directions on NextSeq 500. The sequenced raw data of the JB150 strain was filtered to obtain high quality reads using Trimmomatic *v0.35*. Complete nucleotide sequence of the *Streptomyces* sp. JB150 chromosome has been deposited at NCBI under the accession number CP049780.

### Genome annotation and comparative analysis

Genome annotation was performed using Prokka *v1.12* program [25] and NCBI Prokaryotic Genome Annotation Pipeline (PGAP) [26–27]. For genome and proteome comparisons, *Streptomyces* sp. JB150 (present work), *Actinoalloteichus spitiensis* RMV-1378 [10], *Geodermatophilus africanus* DSM 45422 [11], *Jiangella gansuensis* DSM 44835 [12], *Lechevalieria deserti* DSM 45480 [13], *Modestobacter excelsi* 1G6 DSM 107535 [14], *Streptomyces* sp. DH-12 [15] and *Streptomyces* sp. Wb2n-11 [16] were selected. The complete genomes of these strains available in GenBank [27] were downloaded. A bi-directional BLAST analysis was performed using Pathosystems Resource Integration Center (PATRIC *v 3.6.5*) (https://www.patricbrc.org) [28–29] proteome comparison tool using FIGFams for the selected strains. For comparison of the gene families, PATRIC uses a Pearson pairwise average linkage correlation. Other genomic comparisons and data analyses were performed using the online tools, BLAST available at the NCBI and Sanger web sites (www.ncbi.nlm.nih.gov/BLAST/ and http://www.sanger.ac.uk/), Kyoto Encyclopedia of Genes and Genomes (www.genome.jp/kegg/), and ModelSEED (http://modelseed.org/) for metabolic pathway analysis. Genes that were annotated as transporters were selected and predictions were made by performing a BLAST search against sequences that were downloaded from the TransportDB (Elbourn *et al.* 2017). Prediction of BGCs involved in secondary metabolites synthesis was performed using the antiSMASH *v 5.1.2* software [30]. Main features including distribution of putative coding sequences (CDSs), and BGCs, the direction of transcription (+ strand, upper line; - strand, lower line) as well as a circular diagram of the genome were generated using PATRIC. The locus of the replication origin (*oriC*) was predicted using Ori-Finder [31] available at http://tubic.tju.edu.cn/Ori-Finder2/. The protein families were clustered using a web platform OrthoVenn Analysis Software (http://www.bioinfogenome.net/OrthoVenn/) for comparison and annotation of orthologous gene clusters among multiple species [32].

## Results and Discussion

### Genome assembly and gene predictions

The isolate JB150 was recovered from the soil sample collected at the Bikaner region of the arid Thar desert in India. Based on the phylogenetic analysis, the isolate was identified as *Streptomyces* (MT649861), with the closest 16S rDNA sequence identity (98.02%) to that of the type strain *Streptomyces coelescens* strain AS 4.1594 (Fig.1A). The genome sequence of *Streptomyces* sp. JB150 is linear with 72.72% of GC content and has a size of 7.32 Mbp (Table 1). For comparison, 7 other complete genomes of the desert actinomycetes namely *Actinoalloteichus spitiensis* RMV-1378 [10], *Geodermatophilus africanus* DSM 45422 [11], *Jiangella gansuensis* DSM 44835 [12], *Lechevalieria deserti* DSM 45480 [13], *Modestobacter excelsi* 1G6 DSM 107535 [14], *Streptomyces* sp. DH-12 [15] and *Streptomyces* sp. Wb2n-11 [16], with a genome size ranging from 3.15 to 9.53, were also included in the present study. The genome characteristics and information about the genomes have been furnished in Table 1. The annotation and gene prediction were made with the *R*apid *A*nnotations using *S*ubsystems *T*echnology (RAST) and supported with manual annotations based on domain and motif searches (Table S1a, Supporting Information). Further analysis of the annotated genomes was performed with *P*athosystems *R*esource *I*ntegration *C*enter (PATRIC) available at https://www.patricbrc.org (Table S1b, Supporting Information). Among all the analyzed genomes, *L. deserti* has the largest genome (9.52Mbp) while *A. spitiensis*, a cold desert isolate, has the smallest genome (3.14Mbp). Furthermore, the lowest number of 1874 coding sequences (CDSs) in *A. spitiensis* reflects that it requires a relatively fewer number of genes to endure the cold desert environment (Table 1).

**Fig. 1.**
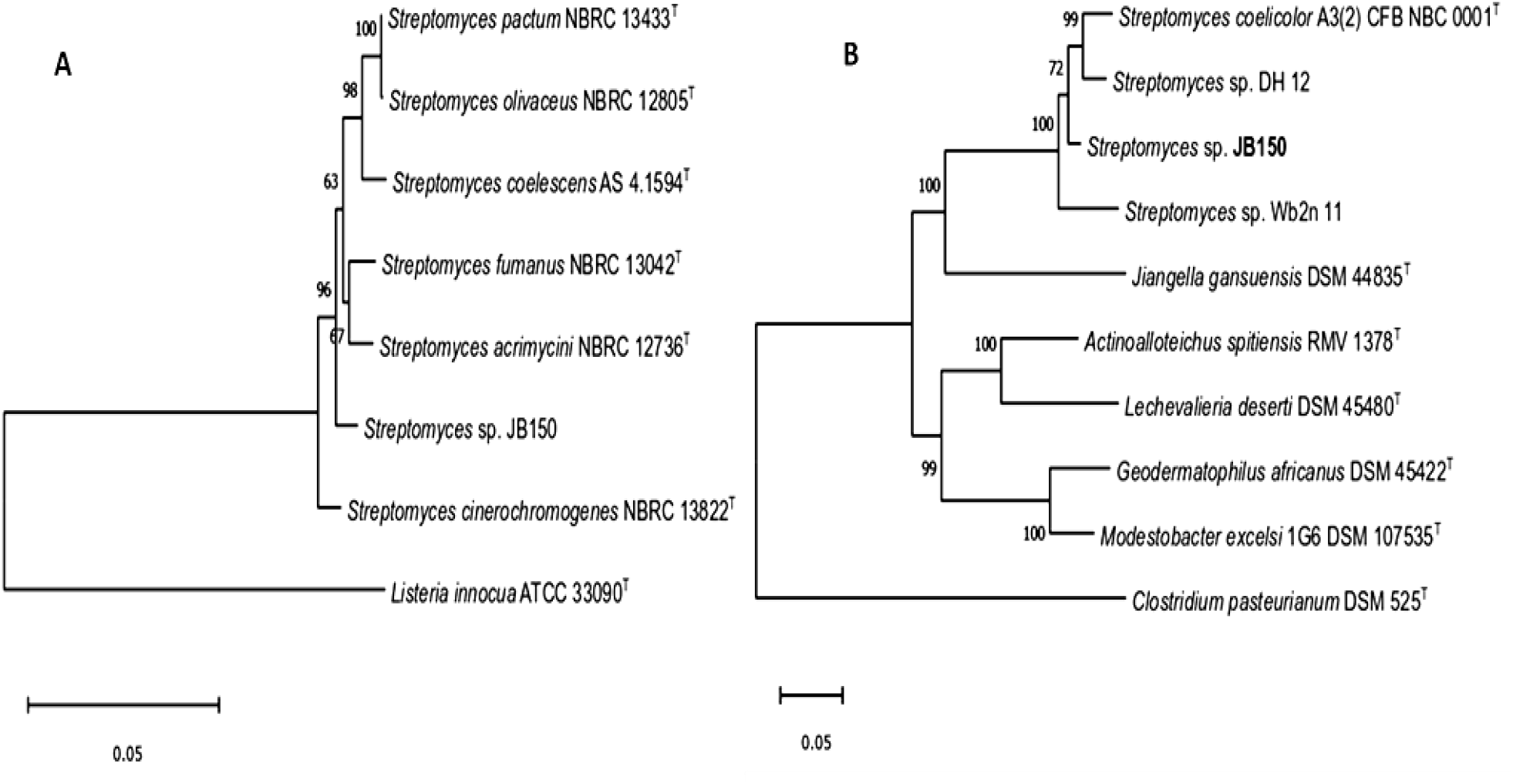
Molecular phylogenetic analysis of conserved 16S rRNA gene fragment of *Streptomyces* sp. JB150. **A**. and *gyrA, rpoB, rsmH* (3 concatenated proteins) of *Streptomyces* sp. JB150 and other seven analyzed genomes of *A. spitiensis*, *G. africanus*, *J. gansuensis, L. deserti*, *M. excelsi*, *Streptomyces* sp. DH-12, and *Streptomyces* sp. Wb2n-11. **B.** by the Neighbor-Joining method. The evolutionary distances were computed using the Kimura 2-parameter method and are presented in the units of the number of base substitutions per site. Evolutionary analyses were conducted in MEGA X.

**Table 1.** General features of the analyzed genomes

| Genome ID | Genome Name | NCBI Taxon ID | Genome Status | Length (bp) | Contigs | GC Content | CDS | Isolation Source |
| --- | --- | --- | --- | --- | --- | --- | --- | --- |
| 1883.1823 | *Streptomyces* sp. JB150 | 1883 | WGS | 7318937 | 1 | 72.73 | 6851 | Thar desert soil, India |
| 1114962.3 | *Actinoalloteichus spitiensis* RMV-1378 | 1114962 | WGS | 3146938 | 6330 | 69.40 | 1874 | Cold desert, Himalaya, India |
| 1860.3 | *Geodermatophilus africanus* DSM 45422 | 1860 | WGS | 5471205 | 52 | 74.30 | 5591 | Sahara desert soil, Ourba |
| 987045.5 | *Jiangella gansuensis* DSM 44835 | 987045 | WGS | 5585780 | 1 | 70.92 | 5343 | Gansu desert soil, China |
| 531939.3 | *Lechevalieria deserti* DSM 45480 | 531939 | WGS | 9529573 | 41 | 68.80 | 9467 | Atacama Desert soil, Chile |
| 2213161.3 | *Modestobacter excelsi* 1G6 DSM 107535 | 2213161 | WGS | 5255906 | 178 | 73.70 | 5275 | Atacama desert soil, Chile |
| 2072509.4 | *Streptomyces* sp. DH-12 | 2072509 | WGS | 7604612 | 13 | 72.80 | 7050 | Thar desert soil, India |
| 1931.11 | *Streptomyces* sp. Wb2n-11 | 1931 | WGS | 8228099 | 5 | 71.03 | 8134 | Sinai desert soil, Egypt |

The genome of *Streptomyces* sp. JB150 was predicted to contain 6851 coding sequences (CDSs), 26 repeat regions, 68 tRNAs, 4 rRNAs, 4669 genes, 263 pseudogenes, and 2182 genes coding for hypothetical proteins. The genome characteristics of *Streptomyces* sp. JB150 has been furnished in Fig. 2.

**Fig. 2.**
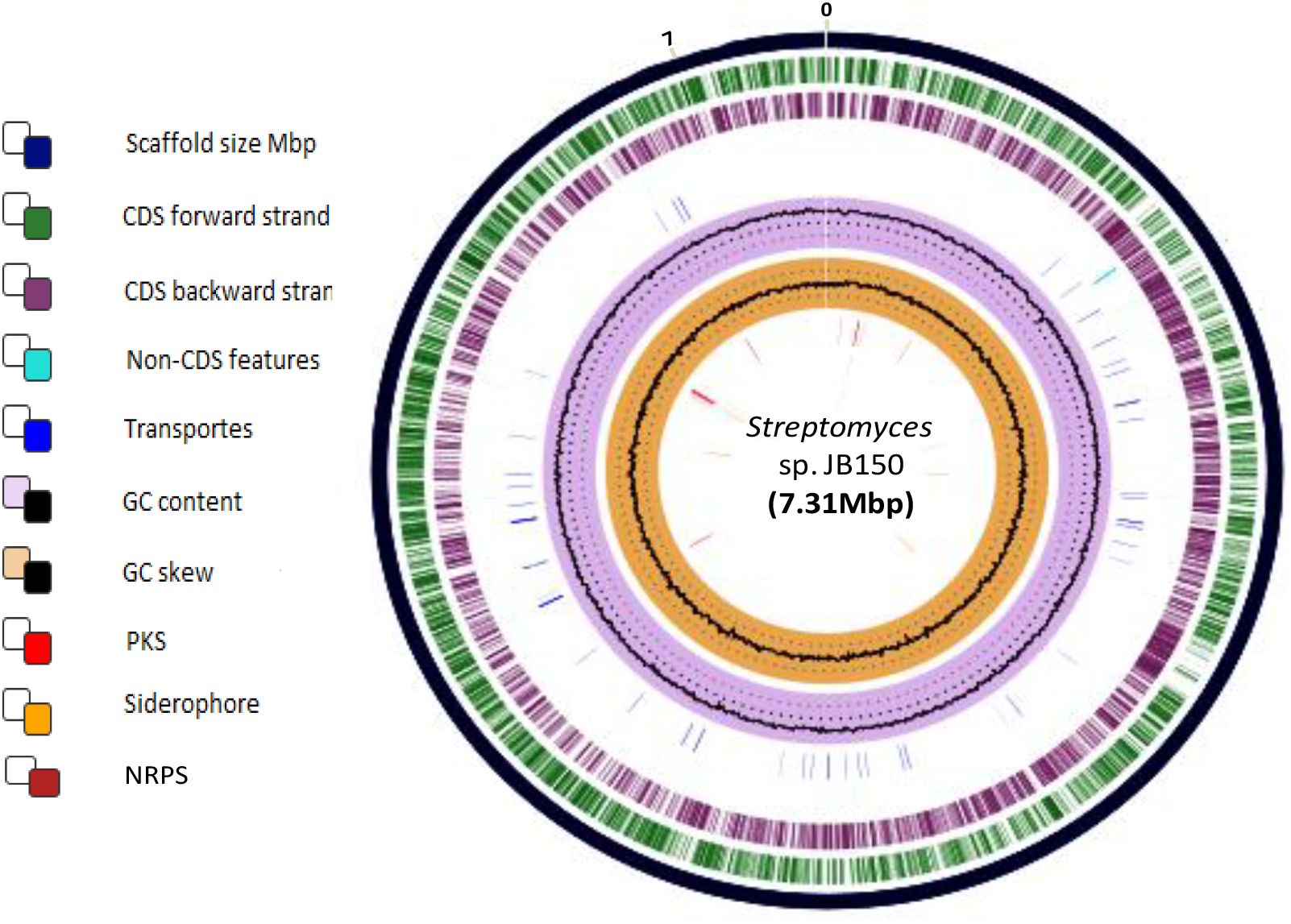
Genome characteristics of *Streptomyces* sp. JB150. The circular map was retrieved from PATRIC (https://patricbrc.org/view/Genome/1883.1823). Colors depict the different classification types of gene clusters along the sequenced genome. The description of each circle is represented from the outermost circle to the innermost. Tick marks representing the predicted CDS on the positive strand and negative strands.

The origin of replication *oriC* is located after the *dnaA* gene (PEG.3617) (Table S1b, Supporting Information) at position 3772751:3773830 (+) of scaffold 1, with the *oriC* repeats spanning over a 1079bp region. Three highly conserved sequences (*gyrA, rpoB, rsmH*) were used to construct a Neighbor-Joining (NJ) phylogenetic tree of all compared genomes, which represents the evolutionary relatedness among the analyzed strains in the present study. An outgroup was formed using the sequence of *Clostridium pasteurianum* DSM 525 (Fig.1B).

### Secondary metabolites

*In silico* genome analyses with the Secondary Metabolite Analysis Shell (antiSMASH *v5.1.2*) algorithm [30], BAGEL [33], PRISM [34], and metabolic mapping using ModelSeed [35] on the *Streptomyces* sp. JB150 genome revealed the presence of 24 biosynthetic gene clusters (BGCs) involved in the production of different secondary metabolites (Fig 3 and Table 2). Many of its gene clusters code for the metabolic compounds that exhibit relative similarity to already known clusters (13% to 100%) except 7 out of 24 clusters (0-6% relative similarity) which possibly encode for new chemical scaffolds. The location of most of the identified clusters resides in both extremes of the scaffold (Fig 3). Out of 24 clusters, 5 BGCs were identified for terpenes in the genome of JB150. Terpenoids are produced by all kingdoms of life including bacteria, fungi, and protists [36–39]. For the synthesis of terpenoids, JB150 was observed to contain terpene synthase genes precursors including farnesyl diphosphate (sesquiterpenes, C15) (EC 2.5.1.10) (PEG.4623) and geranylgeranyl diphosphate (diterpenes, C20) (EC 2.5.1.29) (PEG.126) (Table S1b, Supporting Information).

**Fig. 3.**
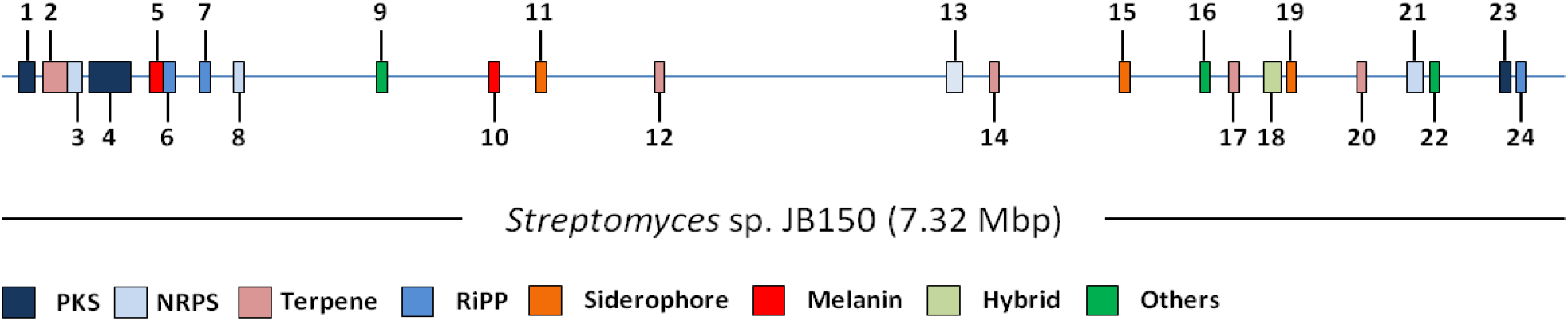
Localization of different secondary metabolite gene clusters in *Streptomyces* sp. JB150

**Table 2.**
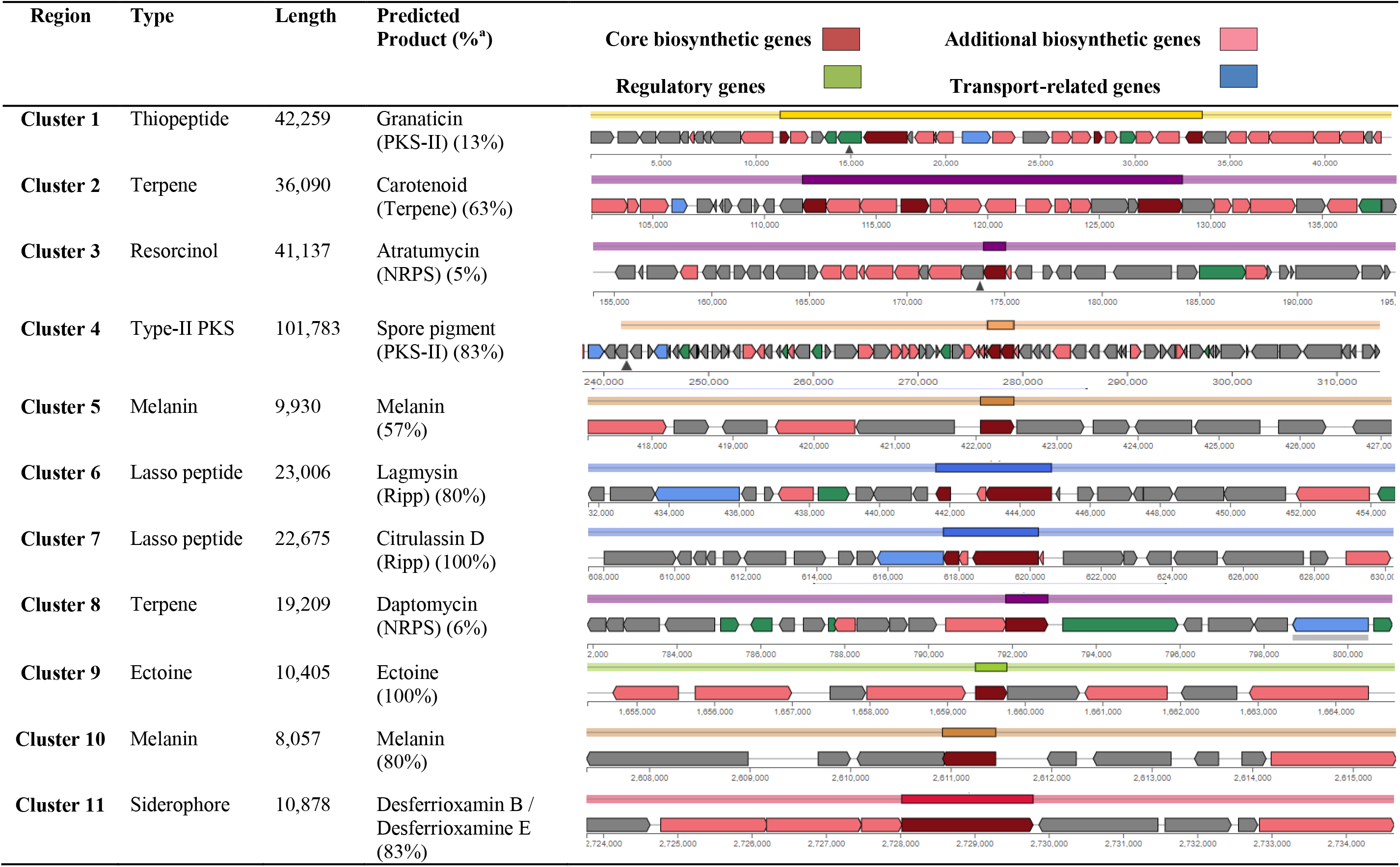
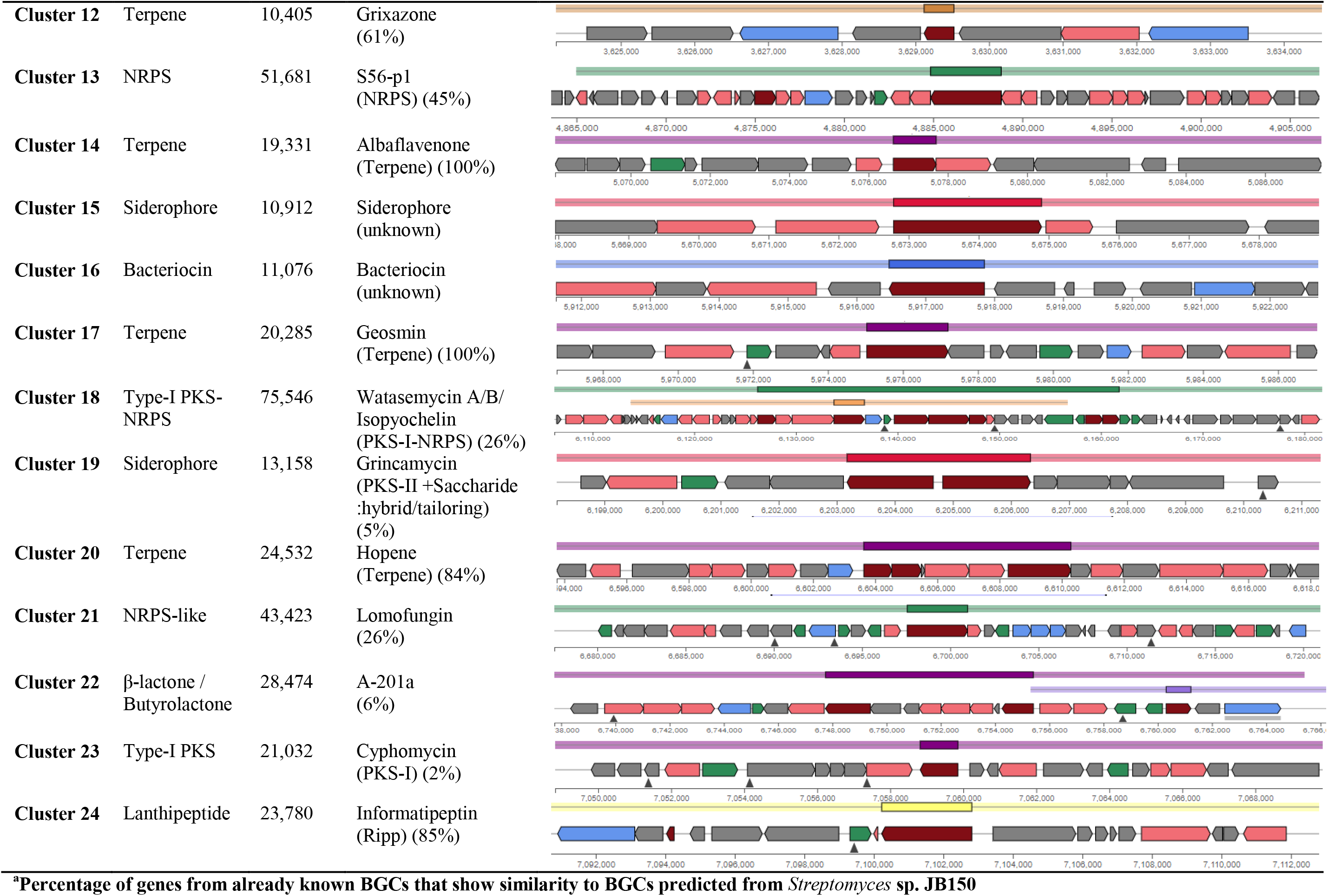
Predicted secondary metabolite gene clusters and their organization in *Streptomyces* sp. JB150 genome

The genome of *Streptomyces* sp. JB150 also encodes two BGCs for melanin that could defend the strain against thermal and biochemical stresses like heavy metals and reactive oxygen species generated by solar UV radiation in the desert environment [40]. The melanin synthesized by the strain could further facilitate its metal ions chelating ability [41] and may help to provide structural rigidity to the cell wall [42]. The genome of JB150 also contains the ectoine gene cluster which may assist the strain to tolerate osmotic stress by accumulating it or de novo synthesizing. Ectoine, 1,4,5,6-tetrahydro-2-methyl-4-pyrimidine carboxylic acid is one of the most commonly found osmolytes in *Streptomyces* [43]. Its osmoprotective function may affect the stability and correct folding of proteins to protect biomolecules such as enzymes and nucleic acids under stress conditions commonly encountered in the desert environment [44–46].

BGCs encoding for secondary metabolites such as atratumycin (cinnamic acid-bearing product) [47], A-201a (nucleoside antibiotic) [48], daptomycin (cyclic lipopeptide produced by a non-ribosomal peptide synthetase) [49], lomofungin (phenazine antibiotic) [50], granaticin (aromatic polyketide) [51], grincamycin (angucycline glycosides) [52], grixazone (parasiticide) [53], watasemycin (iron-chelating NRP) [54], lagmysin (class II lasso peptide) [55], and S56-p1 (contains a unique hydrazine moiety) [56] were appeared to be JB150 specific. The structure of these compounds was predicted using Chemspider and PubChem and has been furnished in Fig 4.

**Fig. 4.**
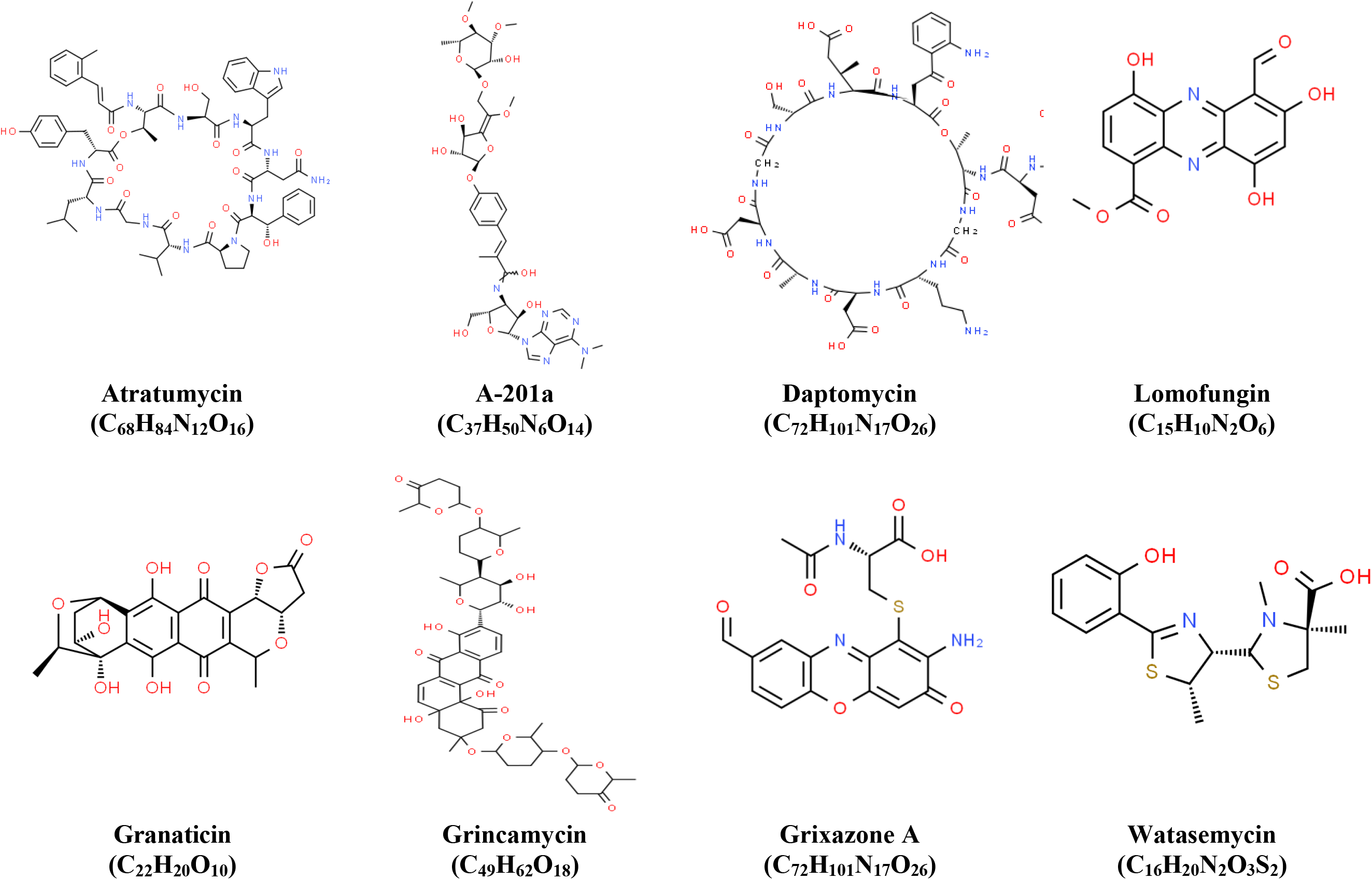
Predicted structures of secondary metabolites unique to *Streptomyces* sp. JB150

**Fig. 5.**
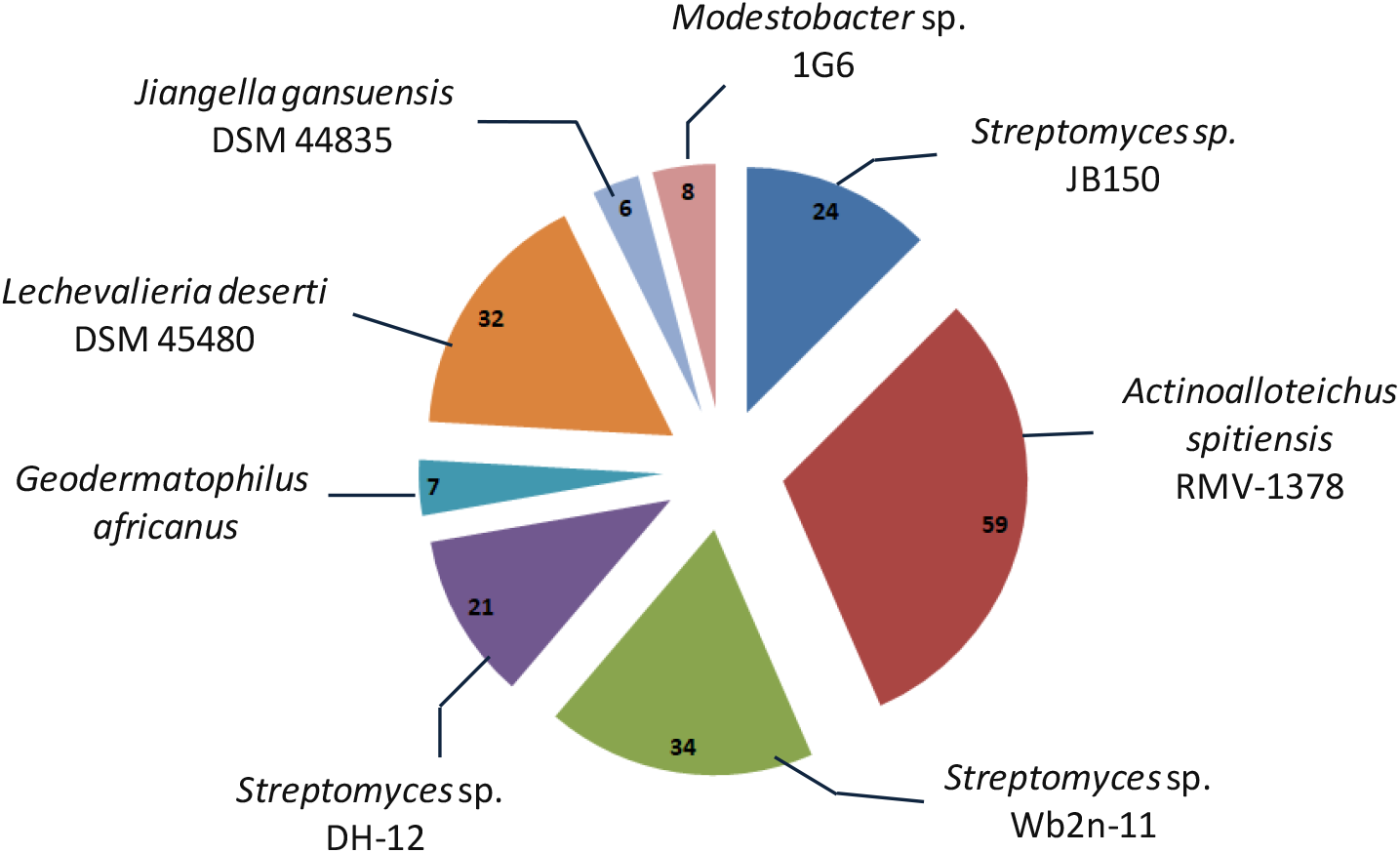
Comparison between the abundance of secondary metabolite gene clusters among *Streptomyces* sp. JB150 and other seven analyzed genomes of *A. spitiensis*, *G. africanus*, *J. gansuensis, L. deserti*, *M. excelsi*, *Streptomyces* sp. DH-12, and *Streptomyces* sp. Wb2n-11

A comparison between the abundance of secondary metabolite gene clusters of distinct types reveals that apart from PKS and NRPS genes, the genomes of JB150 and other 7 analyzed strains have a large number of lanthipeptide and lasso peptide gene clusters (Table S1c, Supporting Information). Lanthipeptide are lanthionine-containing peptides are known as lantibiotics [57] while lasso peptides constitute 15 to 24 amino acids and are ribosomally synthesized and post-translationally modified peptides (RiPPs). RiPPs are characterized by an interlocked structure, the formation of which is dependent on the N-terminal macrolactam ring, and a C-terminal tail [58–61]. Interestingly, despite having the smallest genome among the analyzed strains, the highest number of BGCs was observed in *A. spitiensis* while the genome of *J. gansuensis* was observed to contain only 6 BGCs (Table S1c, Supporting Information). Besides, the prediction of BGCs for secondary metabolites among the 7 genomes using RAST unveils the presence of a unique gene cluster for thiazole-oxazole-modified thiazolemicrocin (TOMM) biosynthesis in JB150. It comprises TOMM dehydrogenase (protein B) (PEG.15) and cyclodehydratase (protein C) / docking scaffold (protein D) (CD fusion protein) (PEG. 18) (Table S1a, Supporting Information). TOMMs are a class of ribosomally synthesized and post-translationally modified peptides (RiPPs), apparently defined by the existence of heterocycles and encompass diverse families, including the linear azol(in)e-containing peptides, cyanobactins, thiopeptides, and bottromycins [62].

### Stress response

The genome of *Streptomyces* sp. JB150 encodes for 62 (4.98%) stress-responsive genes, among which the most abundant genes were appeared to be involved in regulating osmotic stress (9), oxidative stress (18), sigma B mediated stress response (28), detoxification (6), dimethylarginine metabolism (4), periplasmic stress (1), and bacterial hemoglobins (1) (Fig.6). The JB150 strain also appears to produce ectoine as a compatible solute to prevent osmotic stress. For its synthesis and to regulate osmoadaptation, a highly conserved gene cluster *ectABCD* was identified in the genome that encodes L-2,4-diaminobutyric acid acetyltransferase (EctA) (EC 2.3.1.178) (PEG.1639), diaminobutyrate-2-oxoglutarate transaminase (EctB) (EC 2.6.1.76) (PEG.1640), L-ectoine synthase (EctC) (EC 4.2.1.108) (PEG.1641) and ectoine hydroxylase (EctD) (EC 1.14.11.55) (PEG.1642) (Table S1a, Supporting Information). The genome analysis revealed that JB150 has the capacity to synthesize osmoprotectants such as glycine betaine from choline. The enzyme encoded by *betB* (betaine aldehyde dehydrogenase) (EC 1.2.1.8) involved in the glycine betaine biosynthesis pathway was identified. To maintain a positive cellular turgor, a critical parameter for normal metabolic functions, glycine betaine may act as one of the efficient osmolytes. The strain could either assimilate it from the surroundings or synthesize it from choline. Osmotolerance may further be achieved by accumulating osmoprotectants such as proline, biosynthesis of which might be regulated by proline iminopeptidase (EC 3.4.11.5) (PEG.1795) and pyrroline-5-carboxylate reductase (EC 1.5.1.2) (PEG.4077) in JB150 (Table S1a, Supporting Information). Notably, to resist the detrimental effects of high osmolarity environments, the genome analysis of JB150 revealed yet another glycine betaine uptake system encoded by OpuABC operon. It comprises three structural genes: *opuAA* (ATP-binding protein) (EC 3.6.3.32), *opuAB* (permease protein), and *opuAC* (glycine betaine-binding protein) (Table S1b, Supporting Information).

**Fig. 6.**
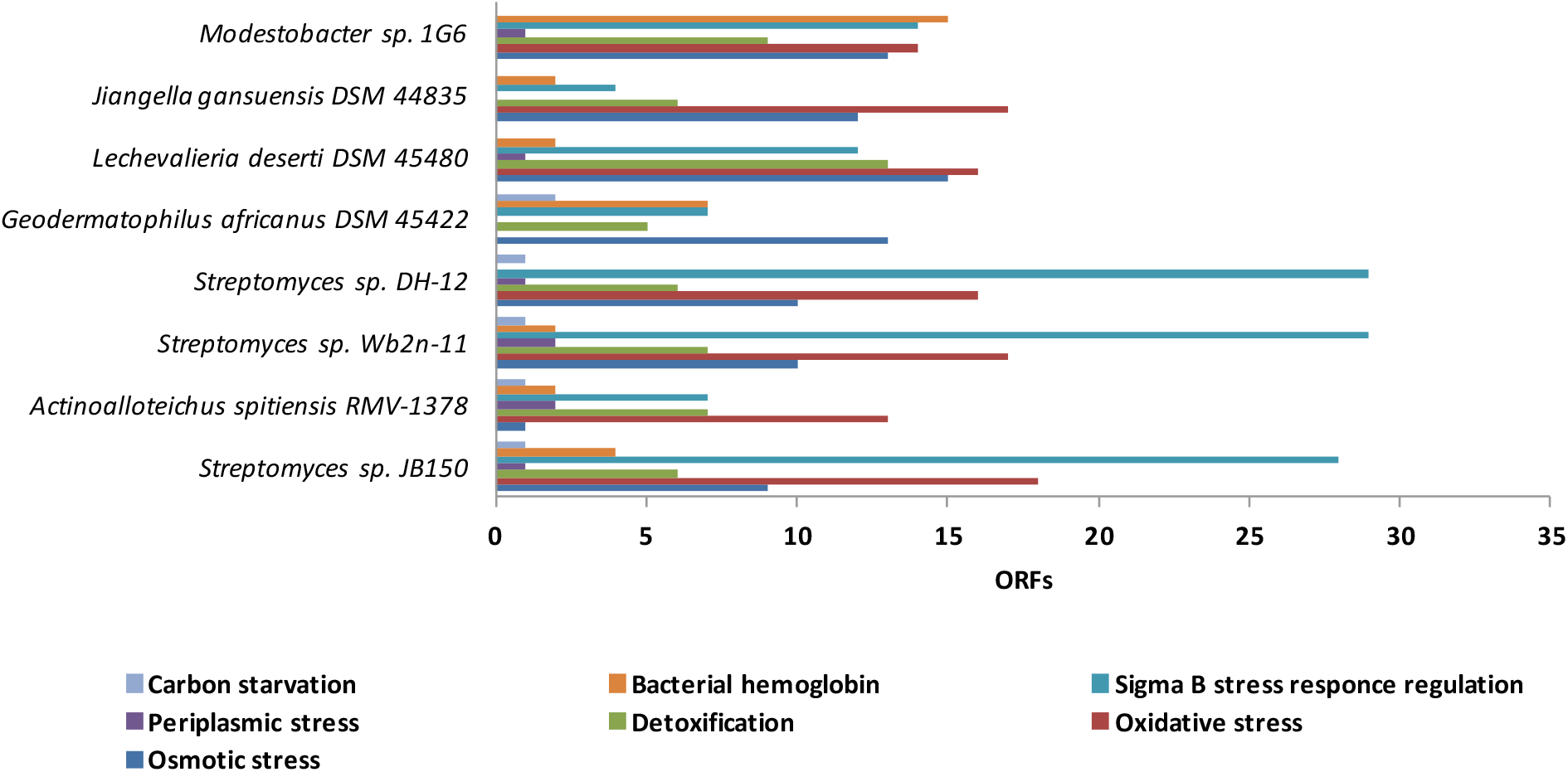
The analysis of **s**tress-responsive genes in the genomes of *Streptomyces* sp. JB150 and other seven analyzed desert-actinomycetes, *A. spitiensis*, *G. africanus*, *J. gansuensis, L. deserti*, *M. excelsi*, *Streptomyces* sp. DH-12, and *Streptomyces* sp. Wb2n-11

Furthermore, for the detoxification of reactive oxygen species (ROS), JB150 is endowed with AhpC (Alkyl hydroperoxide reductase C-like protein) (EC 1.11.1.26) (PEG.2107), thioredoxin peroxidase (EC 1.11.1.15) (PEG.3159), KatE (catalase) (EC 1.11.1.6) (PEG.4828), superoxide dismutase [Fe-Zn] (EC 1.15.1.1) (PEG.2471) and nickel-dependent superoxide dismutase (EC 1.15.1.1) (PEG.4877). The strain JB150 also has regulatory mechanisms for oxidative stress resistance. The proteins OxyR (hydrogen peroxide-inducible genes activator) (PEG.4632), and SoxR (redox-sensitive transcriptional activator/regulator) (PEG.1456) (Table S1a, Supporting Information) were identified to regulate the expression of peroxide and superoxide defense regulons [63–64]. Homologs of OxyR, acting primarily as transcriptional activators, are widespread among actinomycetes [65]. Similarly, the defense mechanisms against oxidative and nitrosative stress could be controlled in JB150 by SoxR and OxyR [66].

According to the genome sequence data from many bacterial species, a diverse number of alternative sigma factors, ranging from one in *Mycoplasma genitalium* [67] sigma factor to about sixty-six different sigma factors in *Streptomyces coelicolor* [68] have been identified. Most alternative sigma factors in actinomycetes belong to the protein family that is homologous to σ^70^ (σ^D^) of *Escherichia coli* [69]. The other known family of σ54 sigma factors in *E. coli* [70] has not been commonly found in actinobacteria [71]. In the present analysis, sigma B (σB; RNA polymerase sigma factor) encoded by the *sigB* operon, a stress-responsive alternative sigma factor, was identified in JB150. The operon includes seven additional genes, which encode RsbT (co-antagonist protein RsbRA) (PEG.215), RsbS, (negative regulator of sigma-B) (PEG.216), RsbT (anti-sigma B factor) (PEG.217), RsbX (phosphoserine phosphatase) (PEG.218) and a serine phosphatase RsbU (regulator of sigma subunit) (PEG.219). The presence of alternative σ^B^ in the genome could contribute to the survival of JB150 under the extreme conditions of starvation and osmotic imbalance [72]. Moreover, to sustain under oxygen limitation or oxidative and nitrosative stress, the strain JB150 also possesses nitric oxide dioxygenase (flavohemoglobin) (EC 1.14.12.17) (PEG.6734) and HbO (hemoglobin-like protein) (PEG.2538) (Table S1b, Supporting Information). The expression of many flavohemoglobin genes might be induced under adverse environmental conditions such as nitrosative stress or oxygen deprivation. In JB150, several chaperones were also identified that may play their roles in the transit of outer membrane proteins through the periplasm [73]. Among them, PpiB (survival protein precursor; peptidyl-prolyl cis-trans isomerase) (EC 5.2.1.8) (PEG.6771), and DegP (HtrA protease/chaperone) (PEG.3536) proteins were observed to be prominent. A gene *romA* (PEG.5987) that pleiotropically inhibits the expression of outer membrane proteins (OMPs) was also identified in the JB150 genome. The regulatory components of σE, RseA (negative regulatory protein) (PEG.4758), and RseP (an intramembrane protease) and RasP/YluC (implicated in the cell division based on FtsL cleavage) (PEG.5262) were also identified in JB150 (Table S1b, Supporting Information).

Comparison between JB150 and 7 other analyzed genomes revealed the highest number of stress-responsive genes (65) in *Streptomyces* sp. Wb2n-11 while *A. spitiensis* contains only 28 number of these genes (Fig. 6 and Table S3, Supporting Information). The small number of these genes in *A. spitiensis* suggests that the strains native to the hot deserts employ more number of genes to exert stress-response for their survival under high temperature than the strains native to the cold deserts. The genes which are commonly shared by all the analyzed strains belong to choline/betaine synthesis, desiccation, glutathione synthesis, and glutathione related non-redox reactions. JB150 and in the other two analyzed *Streptomyces* genomes, a high number of genes (>28) implicated in σB stress response regulation were also identified. *G. africanus* and *M. excelsi* strains possess 7 and 15 genes of bacterial hemoglobins respectively while in *L. deserti*, 15 numbers of genes were more likely to be involved in regulating osmotic stress genes (Fig. 6 and Table S3, Supporting Information). Collectively, all these genetic attributes could confer these strains many potential advantages to thrive in the desert ecosystems.

### Membrane Transport

Transport proteins have an indispensable role in mediating communication between the intracellular and the extracellular environment [74]. They also determine the synthesis and export of the secondary metabolites [75]. Therefore, a study was performed in the present work using TransportDB 2.0 (http://www.membranetransport.org/) [76] to determine the presence of various transporters in JB150 and other analyzed genomes. In the JB150 genome, a total of 591 membrane transporter families were identified. The strain possesses a rich repertoire of ATP-binding cassette (ABC) (56.35%) and major facilitator superfamily (MFS) (14.04%) transporters (Fig.7 and Table S3, Supporting Information). ABC transporters may assist the strain in the translocation of ions, sugars, amino acids, vitamins, lipids, antibiotics, drugs, larger molecules such as oligosaccharides, oligopeptides, and high molecular weight proteins [77]. Similarly, MFS transporters might play their role in the transport of secondary metabolites including antibiotics [78]. Other than these, the transporters belong to amino acid-polyamine-organocation (APC) family, drug/metabolite transporter (DMT) superfamily, and solute: sodium symporter (SSS) family were also identified in high numbers. The APC superfamily of transporters is mainly concerned with the uptake of amino acids and their derivatives while DMT transporters are primarily concerned with the transport of activated sugars for glycolipid and polysaccharide synthesis [79]. Having 9 ORFs of the SSS family of solute:Na^+^ symporters, a constituent member of the APC superfamily [80], JB150 could transport a wide variety of solutes such as short monocarboxylic acids, sugars, etc.

**Fig. 7.**
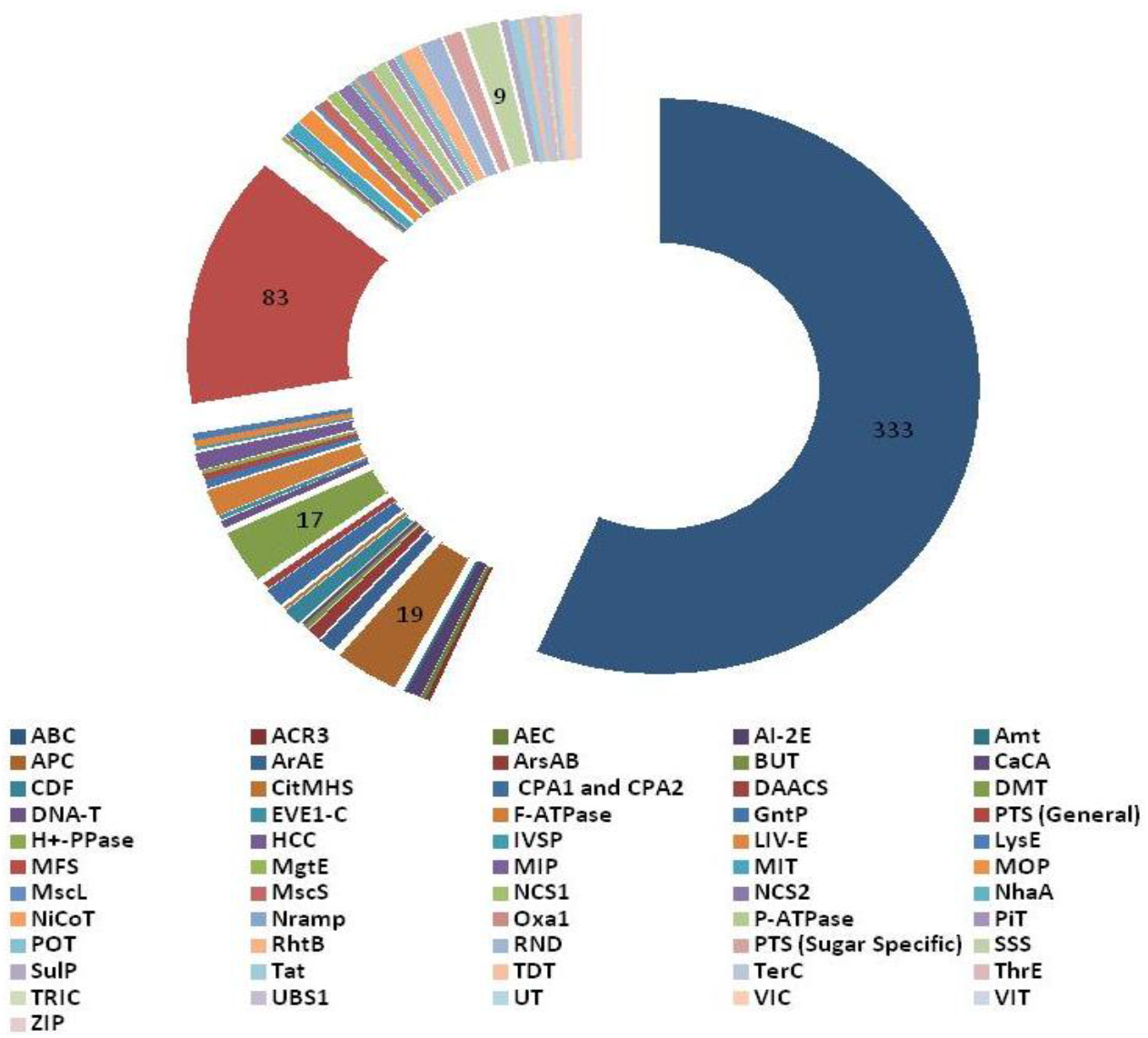
Predicted membrane transporter families in *Streptomyces* sp. JB150 genome

We also observed that the strain JB150 is enriched in H+- or Na+-translocating F-type, V-type, and A-type ATPase (F-ATPase), monovalent cation:proton antiporter-1 (CPA1/CPA2) and resistance-nodulation-cell division (RND) transporters. The transporter analysis also showed that JB150 possesses, MscS and MscL (large and small conductance mechanosensitive ion channel) transporters which may seem to assist the strain to reduce turgor pressure by releasing cytoplasmic solutes [81]. JB150 also contains twin-arginine targeting (Tat) family of transporters which are appeared to involve in nutrient absorption. These transporter families seem to play a prominent role in nutrient acquisition from complex sources, permitting growth when soluble nutrients are not readily available, particularly in the desert environment. Notably, the Tat substrates have been identified in *S. coelicolor,* which constitute a variety of hydrolytic enzymes [82]. Many other transporter families were also identified in JB150, appeared to be involved in taking up nutrients, which include H^+^ symporter (GntP), dicarboxylate/amino acid: cation (Na^+^ or H^+^) symporter (DAACS), proton-dependent oligopeptide transporter (POT), and ammonia transporter channel (Amt) (Fig.7 and Table S3, Supporting Information). These transporters could enable JB150 to make use of alternative carbon and nitrogen sources to obtain energy and thereby increase its competitiveness in response to ecological stresses [83–86]. Additionally, the multiple sugar ABC transporter, general and sugar specific PTS may further assist JB150 to take full advantage of nutrients under oligotrophic nutrient conditions.

In addition to the above transporter families, JB150 also possesses the transporters to participate in metalloid transport and pH regulation. Transporter families of resistance-nodulation-cell division (RND), multidrug/oligosaccharidyl-lipid/polysaccharides (MOPs), and resistance to homoserine/threonine (RhtB) were also identified, which may be involved in drug/metabolite or homoserine lactone efflux [87]. Furthermore, the members of the threonine/serine exporter (ThrE) could allow JB150 to export threonine and serine under the conditions when the higher concentrations of these substances inside the cell may affect growth [88]. The presence of the metal ion (Mn^2+^) transporter (Nramp) in JB150 reflects its adaptability to reduce Mn^2+^ concentrations [89], the optimal concentration of which is crucial for maintaining the normal physiological function of the cell [90]. Members of the arsenical resistance-3 (ACR3) and arsenite-antimonite (ArsAB) efflux family pump As(III) and Sb(III) might also play their important roles in reducing toxicity in the cells and enhancing survivability [91–93]. Additionally, JB150 also possesses Na^+^:H^+^ antiporters (NhaA) to remove excessive Na^+^ to prevent toxicity and to maintain the pH homeostasis in the cells [94] (Fig.7 and Table S3, Supporting Information).

An examination of the integral membrane constituents of ABC transporters in JB150 and other 7 analyzed genomes revealed that *J. gansuensis* contains a very high number of ABC oligopeptides (13) and ABC dipeptide transporter (11) genes (Table S3, Supporting Information). A high number of ABC peptide transporters in the strain could play crucial roles in the nutrient uptake and signaling processes [95–96]. The peptide transporters specific to di and tripeptides (periplasmic dipeptide-binding ABC transporters) such as DppA, DppB, DppC, DppD, DppF or oligopeptides containing five or more residues (periplasmic oligopeptide-binding protein) such as OppA, OppB, OppC, OppD, OppF were observed to be encoded by genes grouped in operons in the genomes of the analyzed strains (Table S3, Supporting Information). Compared to other analyzed genomes, in the genome of JB150, a relatively high number of cation transporters genes (23) were detected. Copper is important for the cells and considered to be a critical cofactor for proteins such as superoxide dismutase, oxidase, oxygenases, and cytochrome C. However, the presence of excess copper could be toxic and may lead to oxidative damage [97–99]. The presence of 10 copper transport genes and CopCD fusion protein (4) suggest that the strain JB150 is copper hypersensitive, wherein CopD imports copper and CopC facilitates CopD-mediated copper import [100]. Besides, the CopCD fusion protein in JB150 could bind to the copper-dependent regulator CsoR, to regulate the expression of the *copZA* genes encoding a cytoplasmic copper efflux system that pumps copper to the extracellular matrix [101].

We also identified the NikABCDE, CbiMNQO, and NikMNQO ABC-type transporters and secondary transporters belonging to the NiCoT, HupE/UreJ, and UreH families in the genome of JB150 which may facilitate the transport of nickel and cobalt. The transition metals nickel and cobalt serve as essential cofactors for several enzymes involved in a variety of metabolic processes [102–103]. To translocate folded proteins, the twin-arginine dependent translocation pathway or Tat pathway was also identified in JB150. Four proteins TatA, TatB, TatC (TatABC polycistronic operon), and TatE (*tatE* is located separately on the chromosome) were identified which are likely to involve in Tat-dependent translocation [104–106]. The analyzed genomes of *Streptomyces* sp. DH-12, *Streptomyces* sp. Wb2n-11, and *A. spitiensis* contain 4-4 copies of the genes implicated in the Tat pathway (Table S3, Supporting Information). The identification of the multiple Na+/H+ antiporters (14) in *A. spitiensis* genome appears to exert appropriate responses under adverse nutrient conditions and other environmental stresses [107–110]. The gene cluster that encodes a Na+/H+ antiporter [111] comprises seven genes (*mrpABCDEFG*) which may express in response to the stress associated with alkaline pH and sodium ions [109]. In JB150 and other analyzed genomes, the stress associated with alkaline pH and sodium ions might be regulated by Na^+^/H^+^ antiport activity mediated by NhaA, NhaD, and sodium-dependent phosphate transporters (monovalent cation–proton antiporter superfamily) [108]. Furthermore, 8 genes for ECF (energy coupling factor) class of transporters were also identified in JB150. These are known as a family of primary active membrane transporters involved in the uptake of essential micronutrients, such as vitamins and trace metals [112]. The genome of JB150 also possesses putative components of type V (PEG.4218) and type VII (PEG.3648, PEG.4693, PEG.5291, PEG.5293, PEG.5295, PEG.5296) secretion systems. The comparison between JB150 and other 7 analyzed genomes revealed that *M. excelsi* possesses the highest number of ECF class transporter genes (11) while these transporters were observed to be absent in *A. spitiensis, Streptomyces* sp. DH-12, *Streptomyces* sp. Wb2n-11, and *L. deserti*. The genomes of *G. africanus*, *J. gansuensis*, *L. deserti*, *Streptomyces* sp. Wb2n-11, and *A. spitiensis* possess the genetic determinants to encode TRAP (tripartite ATP-independent periplasmic) transporters. The system constitutes protein families of DctP, DctQ, and DctM which might assist these strains in the translocation of the solute across the cytoplasmic membrane [113–114] (Table S3, Supporting Information).

### Nitrogen metabolism

Genomic analysis of the JB150 strain using the ModelSEED database (based on KEGG mapping) suggested that the genome of *Streptomyces* sp. JB150 possesses 26 genes of glutamine, glutamate, aspartate, and asparagines (Table S3, Supporting Information). Glutamate and glutamine are the main entry points in the nitrogen assimilation pathway. In JB150, to catalyze the amidation of glutamate to yield glutamine, two distinct glutamine synthetase type I (EC 6.3.1.2) (PEG.1938) and glutamine synthetase type II (PEG.1962) enzymes [115] were observed. A highly conserved operon was also apparent encoding the large and small subunit of glutamate synthases (*gltB*; PEG.1835 and *gltD*; PEG.1836) (Table S1a, Supporting Information). Through the concerted action of the glutamate synthases, ammonium could be incorporated into 2-oxoglutarate, a key intermediate in the tricarboxylic acid (TCA) cycle. A third enzyme, NAD-specific glutamate dehydrogenase (EC 1.4.1.2) (PEG.2915) was also identified in the genome that could catalyze the reductive amination of 2-oxoglutarate to yield glutamate, and is regulated by nitrogen metabolism regulator (GlnR; PEG.3366). GlnR in turn is up-regulated under nitrogen limitation and down-regulated under nitrogen excess [116–117]. In JB150, aspartate aminotransferase (EC 2.6.1.1) (PEG.3721) encoded by *aspC* (regulated by GlnR) was also observed that transfers the amine group from glutamate to oxaloacetate to generate aspartate (Table S1a, Supporting Information).

The analysis of the JB150 genome assisted in the identification of genes that may contribute to the biosynthesis of asparagine. AsnB family represented by asparagine synthetase (glutamine-hydrolyzing) (EC 6.3.5.4) (PEG.5750) was identified. The presence of two asparagine synthetase system in JB150 may ensure sufficient asparagine biosynthesis to confer the strain asparagine prototrophy [118–119]. For the biosynthesis of ammonia and carbamate, the JB150 genome possesses a trimer of a complex formed by three subunits, urease alpha (PEG.898), urease beta (PEG.899) and urease gamma (PEG.900) (EC 3.5.1.5) subunits. For the synthesis of active urease, a complex GTP-dependent process that requires the products of the accessory genes *ureD* (PEG.895), *ureF* (PEG.897) and *ureG* (PEG.896) were also detected. These genes appear to have an important role in urea scavenging, permitting the strain to proliferate and assimilate urea in natural habitats [120]. Similarly, the presence of genes implicated in arginine utilization could confer upon JB150 the ability to efficiently use arginine as a nitrogen source, which is also used in other metabolic purposes such as energy-conservation and polyamine biosynthesis. It seems that its utilization by JB150 is dependent on arginine exporter protein ArgO (PEG.400), arginine deiminase (EC 3.5.3.6) (PEG.5513), arginine beta-hydroxylase (PEG.5528) and arginine decarboxylase (EC 4.1.1.19) (PEG.6823). The pathway regulator arginine biosynthesis repressor ArgR (PEG.1322), a transcriptional regulator of the winged helix-turn-helix (wHTH) family of DNA binding proteins was also identified [121] (Table S1a, Supporting Information).

The genome of JB150 contains a total of 408 genes for amino acid metabolism (Fig.8). In the cysteine and methionine metabolism, JB150 harbors the complete pathways for the transformation of L-cysteine, L-homocysteine, L-methionine, and pyruvate. JB150 also harbors the complete pathways of lysine, valine, leucine, phenylalanine, tyrosine, and isoleucine biosynthesis. Compared to other genomes, JB150 has the highest number of 19 genes for methionine biosynthesis followed by 16 genes for threonine and homoserine biosynthesis (Table S3, Supporting Information). To regulate isoleucine, leucine, valine, β-Hydroxy β-methylglutaryl-CoA (HMG-CoA), and branched-chain amino acid metabolism, 88 genes were identified. Prototrophy for essential amino acids and the capacity to grow without needing external sources can be advantageous to JB150 in poor nutrient soil environments. For putrescine utilization, a complete Puu pathway was also identified in JB150 that metabolizes putrescine via putrescine-binding protein, PotF (ABC transporter) (TC 3.A.1.11.2) (PEG.5234), putrescine transport ATP-binding protein, PotG (PEG.5235) and permease protein PotH (PEG.5236) and PotI (PEG.5237). The presence of the pathway may confer a selective advantage to JB150 because it binds to nucleic acids, stabilizes membrane and stimulates the activity of several enzymes [122–123]. The intracellular deficiency of putrescine may lead to poor cell growth rate and survival in adverse conditions. The comparison of the analyzed genomes revealed that the pathway is likely to be not present in *M. excelsi* and *J. gansuensis* genomes (Table S3, Supporting Information).

**Fig. 8.**
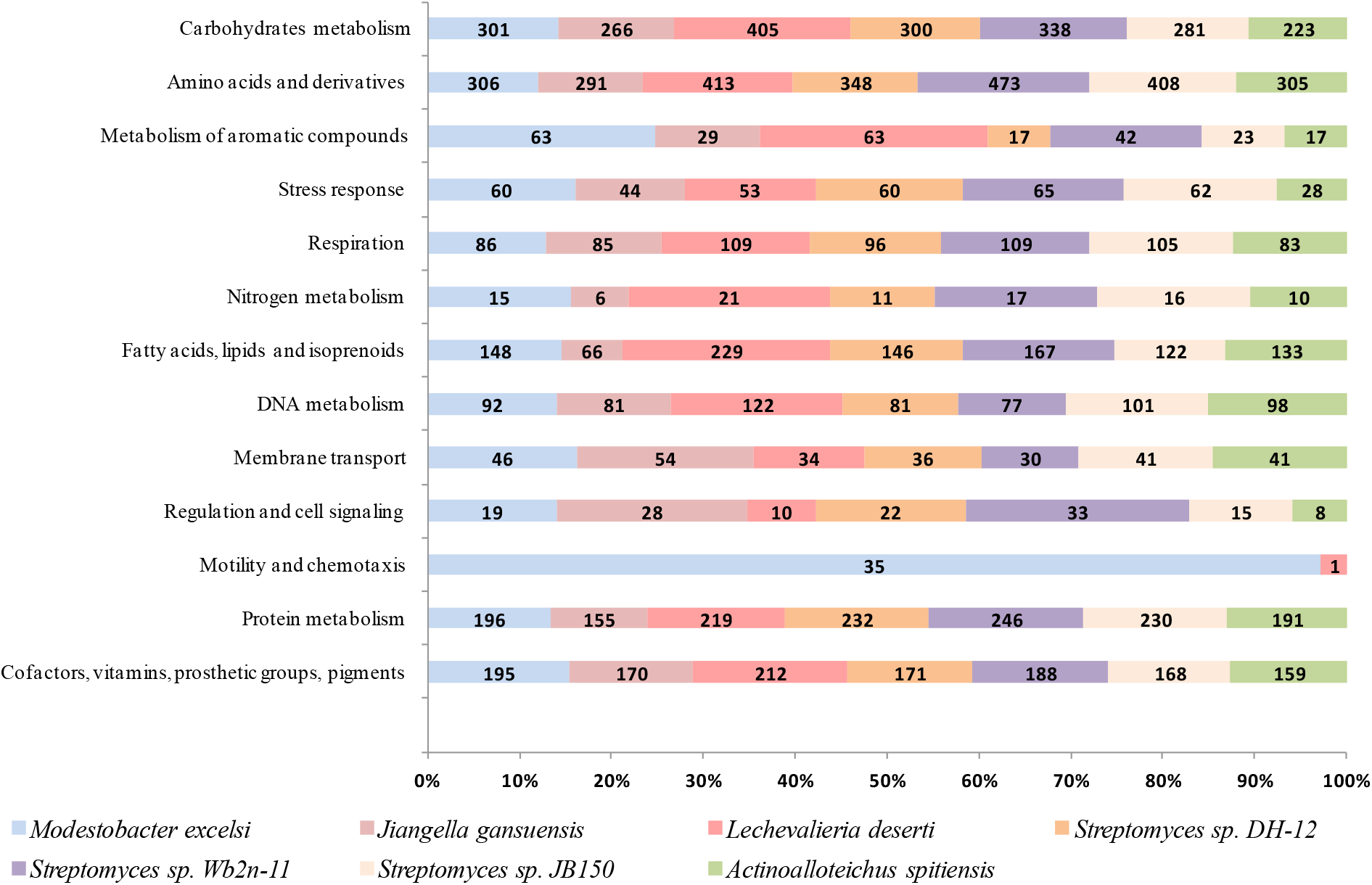
Different subsystem features detected in the genomes of *Streptomyces* sp. JB150 and other seven analyzed desert-actinomycetes, *A. spitiensis*, *G. africanus*, *J. gansuensis*, *L. deserti*, *M. excelsi*, *Streptomyces sp. DH-12*, and *Streptomyces sp. Wb2n-11* using Rapid Annotations using Subsystems Technology (RAST)

In the genome of JB150, the presence of three genes for 1,3-diaminopropane (DAP) synthesis suggest that the strain might be capable of producing small organic polycations polyamines. Biosynthesis of DAP could be achieved in the strain by PLP-dependent enzymes L-2,4-diaminobutyrate decarboxylase (EC 4.1.1.86) (PEG.1829) and diaminobutyrate--2-oxoglutarate aminotransferase (EC 2.6.1.76) (PEG.5375). The DAP pathway was not observed in other analyzed genomes except *Streptomyces* sp. Wb2n-11. A high number of 9 genes encoding 2-keto-3-deoxy-D-arabino-heptulosonate-7-phosphate synthase II (EC 2.5.1.54) (DAHP) (PEG.6) (AroAII homology family) to perform the initial reaction in aromatic amino acid biosynthesis existed in the genome of JB150. The strain may engage DAHP synthase to act as a primary catalyst for phosphoenolpyruvate (PEP) and erythrose 4-phosphate (E4P) to synthesize aromatic amino acids, vitamin K, folic acid, and ubiquinone. The pathway has been known to contribute massively in the biosynthesis of antibiotics, defense metabolites, siderophores, pigments, signaling compounds, and various secondary metabolites [124].

Among JB150 and other 7 analyzed genomes, the highest number of genes (27) for glutamine, glutamate, aspartate, and asparagine biosynthesis was identified in *L. deserti*. It also harbors the maximum number of genes (16), which are likely to be involved in histidine metabolism. An unusually high number of 72 genes seem to be involved in the metabolism of arginine, urea, and polyamines were also observed in the genome of *Streptomyces* Wb2n-11 (Table S3, Supporting Information). In the putrescine utilization pathway, 4 genes were commonly shared by *G. africanus*, *Modestobacter excelsi*, *Streptomyces*. DH-12, *Streptomyces* Wb2n-11, and *Streptomyces* JB150 genomes while the system was found to be missing in *A. spitiensis, J. gansuensis*, and *L. deserti* genomes. The genome of *Streptomyces* Wb2n-11 possesses the highest number of methionine (27) and threonine/homoserine (20) biosynthesis genes. However, the genes for cysteine biosynthesis were present only in JB150, and the other two *Streptomyces* genomes selected in the present analysis. Similarly, a large number of genes (32) for valine degradation were observed only in the genome of *Modestobacter excelsi.* We did not observe the genes for creatine and creatinine degradation in *A. spitiensis*. Notably, the genes for DAP production was identified only in *Streptomyces* Wb2n-11 and JB150 genomes. The genome of JB150 also possesses the highest number of genes (9) for the synthesis of aromatic compounds (DAHP synthase to chorismate) and proline utilization. *Streptomyces* Wb2n-11 was observed to contain a large number of genes (23) for the synthesis of tryptophan, PAPA antibiotics, PABA, and 3-hydroxyanthranilate (Table S3, Supporting Information). The biosynthetic capacity to synthesize a wide range of amino acids and other molecules of the analyzed genomes might be advantageous in the highly underprivileged and famine desert environment.

### Clusters of orthologous groups of proteins, carbohydrate-active enzymes and protein families

All coding sequences (CDSs) on the chromosome of JB150 and other 7 analyzed strains were compared using orthologous clustering analysis (http://www.bioinfogenome.net/OrthoVenn/) [32]. The analysis of CDSs on the JB150 chromosome was classified into 5,030 clusters, among which 1030 orthologs (20.48%) were observed to be commonly shared by all the analyzed strains (Fig. 9A). The genomes of *Streptomyces* JB150, *Streptomyces* DH-12, and *Streptomyces* Wb2n-11 were observed to contain 344, 196, 425 ORFs of unique COGs respectively while these three genomes were observed to share 3494 orthologous (Fig. 9B). The highest number of 1481 singletons (strain-specific proteins) were identified in *Streptomyces* Wb2n-11, followed by 875 and 1022 singletons in *Streptomyces* DH-12, and *Streptomyces* JB150 respectively (Fig.9A). *L. deserti* was observed to have the largest number (3145) of strain-specific proteins that can further be attributed to the large size of its genome (9.53 Mbp) while the lowest numbers of strain-specific proteins were apparent in the genome of *Streptomyces* DH-12. We also conducted a COG function category comparison among the selected genomes using Go enrichment analysis which revealed that the core genomes of these strains harbor a large proportion of commonly shared gene clusters which appear to be involved in translation (57 ORFs), regulation of transcription (DNA-templated) (17 ORFs), DNA replication (12 ORFs), glycolysis (11 ORFs), histidine biosynthesis (9 ORFs), quinone binding (9 ORFs), de novo inosine monophosphate biosynthesis (8 ORFs), protoporphyrinogen IX biosynthesis (8 ORFs), ATP synthesis coupled proton transport (7 ORFs), chorismate biosynthesis (7 ORFs), oxidoreductase activity (6 ORFs), terpenoid biosynthesis (6 ORFs), threonine biosynthesis (6 ORFs), transmembrane transport (6 ORFs), RNA binding (5 ORFs), and proteasomal protein catabolism (4 ORFs). Among 1030 protein clusters, commonly shared by JB150 and seven other genomes (Table S1e, Supporting Information), the highest number of 18 and 14 ORFs were identified to be associated with the cellulose catabolic process of cluster 6 (Fig. 9c) and antibiotic biosynthetic process of cluster 16 (Fig. 9d) respectively. The abundance of these genes might play a significant role in conferring upon the strains to gain a competitive advantage in the extreme conditions of the desert by providing protective measures and ensuring protein synthesis in oligotrophic nutrient conditions.

**Fig. 9.**
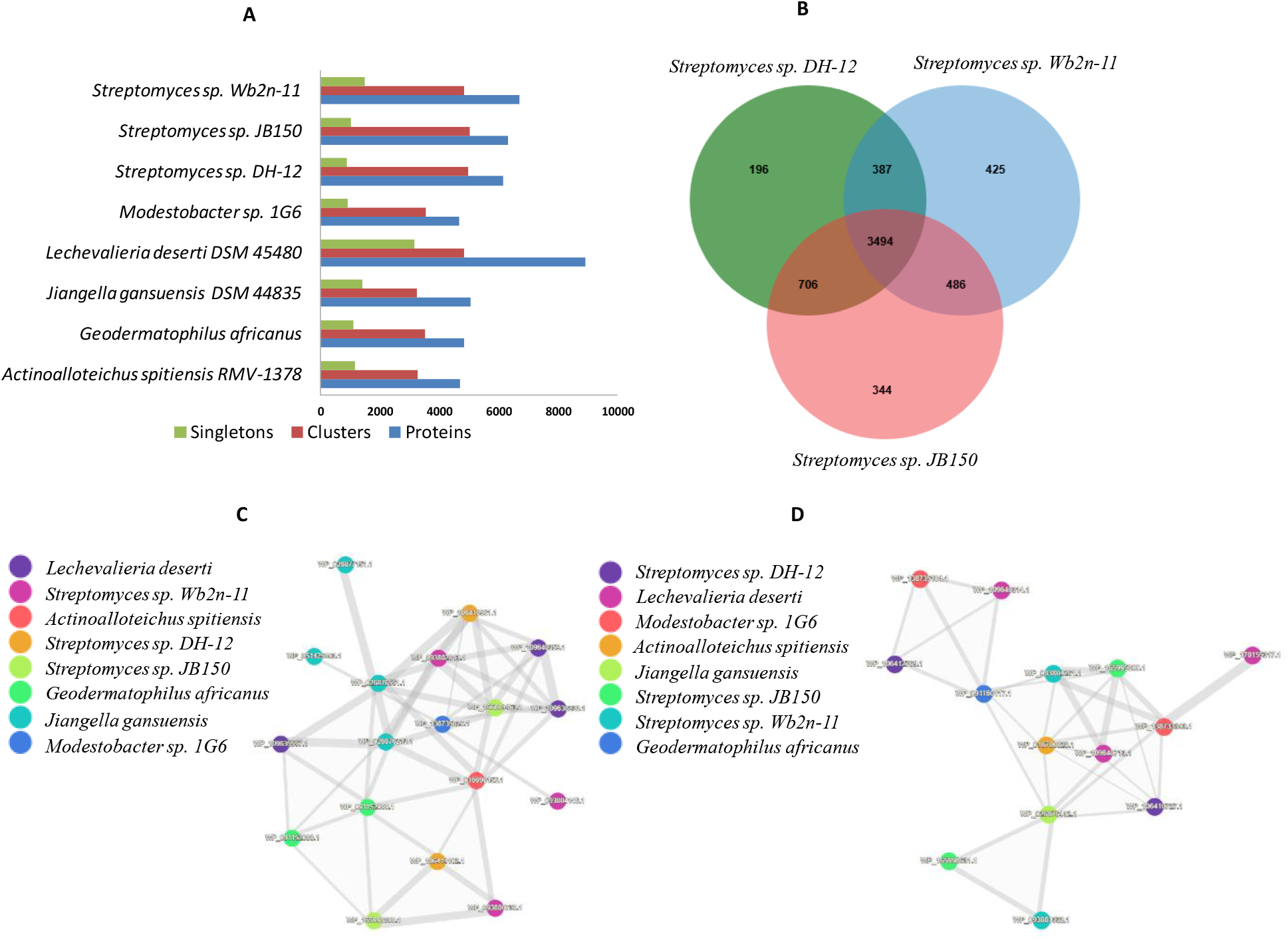
Cluster of orthologous genes (COGs) of proteins in the analyzed genomes. Singletons represent single-copy genes **(A)**, Venn diagram of the number of gene clusters commonly shared by *Streptomyces* sp. JB150 and the other two analyzed *Streptomyces* sp. DH-12, and *Streptomyces* sp. Wb2n-11. The shared core gene number is shown in the center and strain-specific gene number is shown at the corners **(B)**. Predicted protein-protein interactions of clusters involved in the cellulose catabolic process (cluster 6) (GO annotation: 0030245) **(C)**, and antibiotic biosynthetic process (cluster 16) (GO annotation: 0017000) **(D)**. Each node in the above figure is a cluster and the node size represents the proteins in the cluster. Proteins IDs are denoted as WP_ and each edge in the figure represents the relationship between two protein clusters.

To determine the proportions of different functional classes of CAZymes families, the ORFs from JB150 were uploaded on the carbohydrate-active enzyme database (CAZy) [125] and the sequences that match were manually analyzed. The predicted families constitute glycoside hydrolases (GHs), glycosyltransferases (GTs), polysaccharide lyases (PLs), carbohydrate esterases (CEs), auxiliary activity (AA), and carbohydrate-binding functional modules (CBM) (Fig. 10 and Table S1f, Supporting Information). The GHs (EC 3.2.1.X) family of enzymes catalyzes the hydrolysis of glycosidic bonds of carbohydrate substrates in carbohydrate metabolism to degrade starch, cellulose, and hemicellulose [126–127]. We found a total of 128 GHs classified into 46 predicted families in the genome of JB150 (Figure 10 and Table S1f, Supporting Information). GH family classification revealed a total of 22 (17.18%) families with only one GH. In JB150, the GH13 family was observed to be prominently present with 18 genes. The genes in the family include trehalose-6-phosphate hydrolase (EC 3.2.1.93), trehalose synthase (EC 5.4.99.16), malto-oligosyltrehalose trehalohydrolase (EC 3.2.1.141), malto-oligosyltrehalose synthase (EC 5.4.99.15) enzymes seem to be involved in the synthesis of a disaccharide storage reserve trehalose. Intriguingly, among the analyzed genomes, the highest number of trehalose biosynthesis genes (18) was identified in the JB150 genome (Table S3, Supporting Information). Similarly, in CAZyme identification, a total of 12 GT families including 70 GTs were identified. Among these, 5 families were identified as a family of one GT gene. GTs (EC 2.4.X.Y) catalyze glycosyl group transfer to specific acceptor molecules, leading to the formation of a glycoside bond in the process of biosynthesis of polysaccharides, oligosaccharides, and glycoconjugates [128–130]. In JB150, the GT2 and GT4 families were prominently present which possessed 27 and 20 genes respectively.

**Fig. 10.**
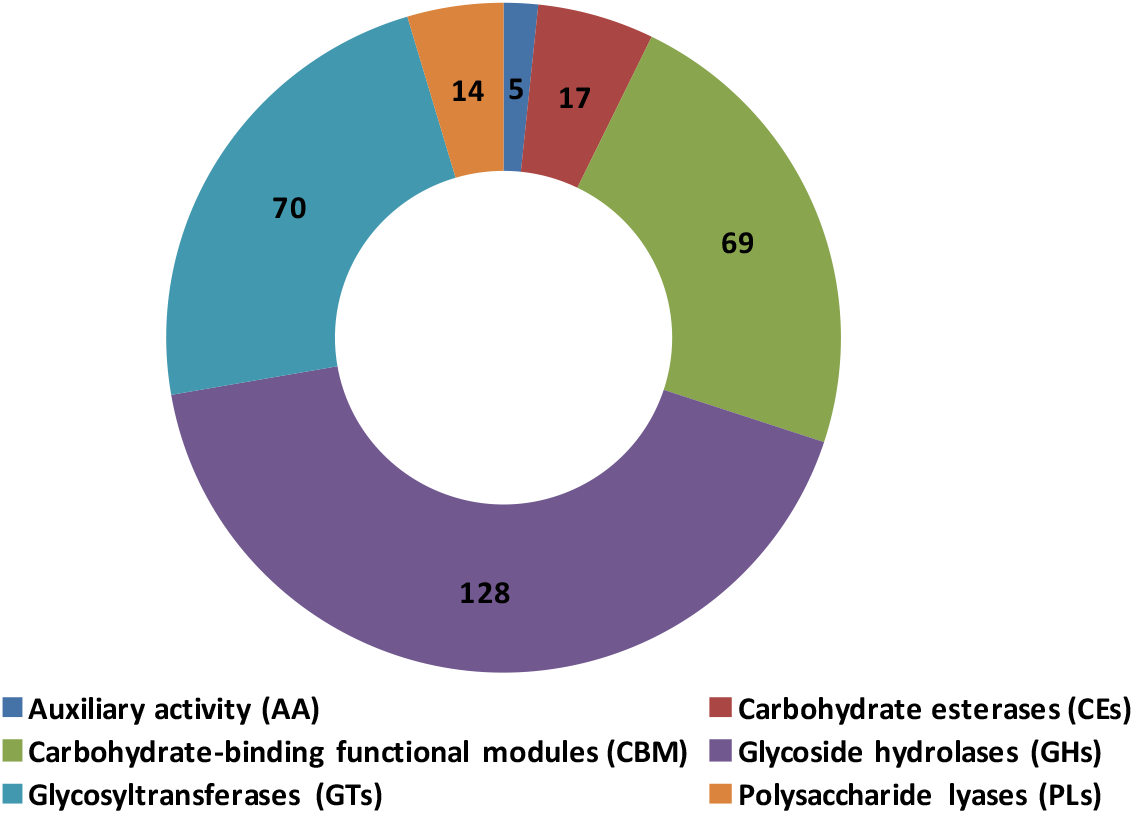
Functional classes of Carbohydrate-Active Enzymes (CAZymes) in the genome of *Streptomyces* sp. JB150

The genome of JB150 contains, a total of 14 PLs (EC 4.2.2.-) classified in 6 families among which PL1 family was observed to be prominently present with 8 genes. It seems that PLs could direct JB150 to act on polysaccharides to produce unsaturated polysaccharides using a beta-elimination mechanism [131]. The diverse members of PL families including PL1 and PL3 in the JB150 genome also suggest that the strain is capable of processing various polysaccharides. A class of esterases known as CEs was also identified in JB150 that may catalyze the O-de- or N-deacylation to remove the ester of substituted saccharides [125]. Similarly, a total of 17 CEs, classified into 7 families were identified wherein CE12 family was assessed to be prominently present with 4 genes (Table S1f, Supporting Information). CEs facilitate the access of GHs to speed up the degradation of substrates in the process of saccharification [132]. The strain also possesses one AA family (AA10) with 05 AAs, which have been originally classified as chitin-binding proteins (CBM33) [133], to catalyze non-carbohydrate structural components. The comparison of JB150 and other seven analyzed genomes revealed a large number of chitin utilization genes (32) in the genome of *L. deserti* (Table S3, Supporting Information). The protein members of this family are currently known as copper-dependent lytic polysaccharide monooxygenases (LPMOs) [134]. In the genome of JB150, a total of 69 CBMs were classified into 15 families among which CBM2 (18 genes) and CBM13 (11 genes) families were observed to be prominent. By utilizing the carbohydrate-binding activity of these enzymes, the catalytic efficiency of CAZymes could be enhanced. However, the hydrolytic activity is not associated with CBMs yet their association with GHs, GTs, and PLs enzymes may lead to the degradation of the substrate more efficiently. Among CBMs, CBM4, CBM5, CBM61, and CBM67 families were observed with only one CBM in the JB150 genome (Table S1f, Supporting Information). CBMs have crucial roles in cellulases and cellobiohydrolases and belong to the GH6 and GH7 families [135].

The proteomes of JB150 and 07 other analyzed strains were also analyzed and compared using the PATRIC PGfams software to identify cross-genus families (CGFs) (Fig 11, Table S2, Supporting Information). From a total of 13143 CGFs, 360 families (7.3%) were commonly shared by JB150 and the other seven analyzed genomes (Table S1g, Supporting Information). Compared to other genomes, JB150 contains 481 unique genes that have no detectable homologs in others. Thus, these genes may be considered as orphan genes (Table S1h, Supporting Information). Among these, the genes for [LysW]-L-2-aminoadipate 6-phosphate reductase (PEG.4676), [LysW]-aminoadipate kinase (PEG.6326), and [LysW]-lysine hydrolase (PEG.4666) were identified in JB150 (Table S1b and S1h, Supporting Information) which could contribute their roles in *α*-aminoadipate (AAA) pathway, a second lysine biosynthesis pathway besides DAP, previously identified in *T. thermophilus* [136–138].

**Fig. 11.**
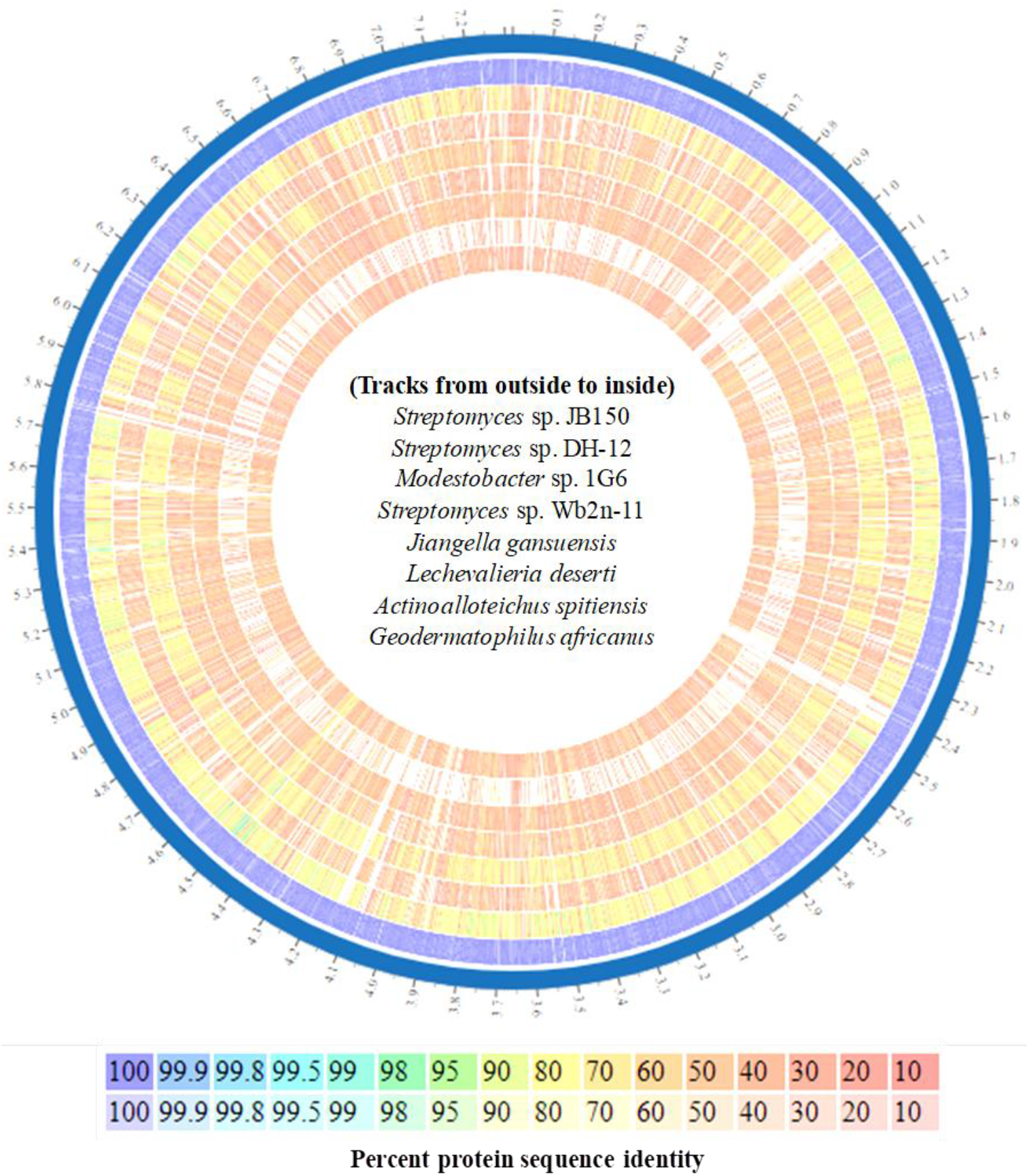
Comparison of proteomes between JB150 and other seven analyzed actinomycetes strains *A. spitiensis*, *G. africanus*, *J. gansuensis, L. deserti*, *M. excelsi*, *Streptomyces* sp. DH-12, and *Streptomyces* sp. Wb2n-11. The genome tracks are from outside to inside as depicted. The exterior (solid-line circle) indicates the scaffolds of the reference genome ±JB150. In each circle, the line indicates one protein homologous to a protein in the reference genome. Percent protein sequence identity exhibits the relationship between proteins.

Another interesting orphan gene cluster that encodes 2,3-diaminopropionate (SbnA; PEG.237), (SbnB; PEG.238), likely to be involved in NRPS-independent siderophore (NIS) synthesis was also observed in JB150. SsgB (PEG.1284), a sporulation-specific cell division protein and SsgG protein (PEG.2832) [139], were also observed in the genome of JB150. Similarly, many orphan genes involved in different pathways were observed to be specific to the JB150 genome include 3-amino-4-hydroxybenzoate synthase (EC 4.1.99.20) (PEG.3487) (grixazone biosynthesis), threonylcarbamoyl-AMP synthase (EC 2.7.7.87) (PEG.4933) (L-threonine metabolism), dUMP forming, deoxycytidine triphosphate deaminase (EC 3.5.4.30) (PEG.6529) (nucleotide metabolism), leucine-specific ABC transporter, substrate-binding protein LivK (TC 3.A.1.4.1) (PEG.6847), peptidyl-prolyl cis-trans isomerase, PpiB (EC 5.2.1.8) (PEG.6771) (proline metabolism), pyridoxal 5-phosphate (PLP)-dependent ornithine decarboxylase (EC 4.1.1.17) (PEG.6525) (polyamines biosynthesis), taurine ABC transporter, substrate-binding protein TauA (PEG.6357), type V secretory pathway (PEG.4219) (adhesin AidA), serine protease (PEG.4696) (putative component of Type VII secretion system), polysaccharide pyruvyl transferase, CsaB (PEG.843) (peptidoglycan-associated polymer biosynthesis), diguanylate cyclase/phosphodiesterase domain 2 (EAL) (PEG.1138) (biosynthesis of secondary messenger, cyclic-di-GMP) etc (Table S1b and S1h, Supporting Information). In JB150, the physiological mechanisms of many of these orphan genes remain to be investigated. The proteome comparisons of the analyzed genomes further revealed that *Streptomyces* JB150 and *Streptomyces* DH-12 share the highest number of orthologues (273) followed by *Streptomyces* Wb2n-11 with which it shared 115 orthologues (Table S1i, Supporting Information). Relative to the other seven analyzed genomes, JB150 includes 132 specialty genes most of which seem to be implicated in conferring on the strain antibiotic resistance and support many transport functions (Table S1j, Supporting Information).

### DNA metabolism

JB150 encodes a complete set of enzymes required for DNA replication, recombination, and DNA repair. The UvrABC proteins (PEG.1746, PEG.1753, PEG.1766) essential to initiate nucleotide excision repair were identified. An excinuclease ABC (subunit A), a paralog of unknown function, was also observed in JB150 (PEG.3877) (Table S1a, Supporting Information). To process and repair double-strand breaks, JB150 possesses a pathway that seems to involve the multifunctional helicase/nuclease complex (RecBCD) (PEG.1425, PEG.2589, PEG.4801). A higher number of genes (20) associated with DNA repair (base-excision) system in JB150 suggest that the strain can efficiently repair damages caused by oxidation, deamination, or alkylation. To initiate this process the system appears to involve DNA glycosylases (formamidopyrimidine-DNA glycosylase) (EC 3.2.2.23) (PEG.6393), G/U mismatch-specific uracil DNA glycosylase (EC 3.2.2.28) (PEG.926), DNA-3-methyladenine glycosylase (EC 3.2.2.20) (PEG.4753) and DNA-3-methyladenine glycosylase II (EC 3.2.2.21) (PEG.1557) to recognize and remove damaged or inconsistent bases. To remove inappropriate bases, endonuclease III (EC 4.2.99.18) (PEG.3929), endonuclease IV (EC 3.1.21.2) (PEG.2753), and endonuclease VIII (PEG.2464) were also identified. To complete the final steps in DNA repair, JB150 also possesses exodeoxyribonuclease III (EC 3.1.11.2) (PEG.3101), exodeoxyribonuclease VII small and large subunits (EC 3.1.11.6) (PEG.4648, PEG4649), DNA polymerase I (EC 2.7.7.7) (PEG.1811), DNA polymerase IV (EC 2.7.7.7) (PEG.1120), error-prone repair homolog of DNA polymerase III alpha subunit (EC 2.7.7.7) (PEG.1503) and DNA ligase (ATP-dependent) (EC 6.5.1.1) (PEG.5966) as well as DNA ligase (NAD(+) dependent) (EC 6.5.1.2) (PEG.5061) [140] (Table S1a, Supporting Information). In the genome of *A. spitiensis*, we identified the highest number of 15 genes of clustered regularly interspaced short palindromic repeats (CRISPER).

The genomes of *Streptomyces* DH-12 and *G. africanus* also bear 9 and 1 genes of CRISPER respectively (Table S3, Supporting Information). It appears that these short repeats confer upon these strains the ability to integrate fragments of foreign DNA spacers into CRISPR cassettes. Consequently, the CRISPR cassettes could be transcribed to synthesize CRISPR RNA to specifically target and slice the genome of the invaders [141–144]. The genomes of *Streptomyces* DH-12 and *G. africanus* encompass the most conserved and abundant *cas* genes and their products namely Cas1, Cas2, Cas3, and Cas4 [145]. These two genomes also contain several specific genes such as *csa1*, *csa2*, *csa3*, and *csa4* of Cse and Csy signature protein families [146]. The system was observed to be missing in *M. excelsi*, *J. gansuensis*, *L. deserti*, *Streptomyces* Wb2n 11, and *Streptomyces* sp. JB150. In the genome of *L. deserti* we identified the highest number of 7 genes (compared to 2 genes in the other analyzed genomes) of type I restriction-modification system which comprises three proteins; one each for restriction (HsdR), modification (HsdM) and DNA binding (HsdS) [147,148] (Table S3, Supporting Information).

### Fatty acids, lipids, and isoprenoids

The search for fatty acid biosynthetic genes in the JB150 genome revealed various putative orthologues to several of the classical fatty acid synthase (FAS) II components. The genome of *Streptomyces* sp. JB150 possesses 122 genes (9.80%) for fatty acid, lipid, and isoprenoid metabolism (Fig.8). The analysis of the annotated genome revealed that many genes encode for the housekeeping enzymes that engage in the saturated fatty acid biosynthesis. These genes lie in a conserved fatty acid biosynthesis (*fab*) cluster that constitutes acyl carrier proteins (ACPs) (PEG.43), (PEG.173) (PEG.285), *fabD* (malonyl CoA-acyl carrier protein transacylase) (EC 2.3.1.39) (PEG.44), *fabF* (3-oxoacyl-[acyl-carrier-protein] synthase, KASII) (EC 2.3.1.179) (PEG.39) and *fabH* (3-oxoacyl-[acyl-carrier-protein] synthase, KASIII) (EC 2.3.1.180) (PEG.38) [149,150] (Table S1a, Supporting Information). JB150 has a single copy of *fabD* gene in the main *fab* cluster and one *fabD* homolog (PEG.2120) suggesting their essential role in fatty acid biosynthesis. Since FabD has been observed to be active with several ACPs [151] the functional role of FabD in JB150 could be predicted to provide malonyl-ACP precursors in both fatty acid and type II polyketide biosynthetic pathways. Similarly, FabH protein which shows broad substrate specificity for short-chain acyl-CoA substrates might have a role in initiating branched and straight-chain fatty acid biosynthesis [152]. Furthermore, the gene catalyzing an essential step in fatty acid biosynthesis encoded by enoyl-[acyl-carrier-protein] reductase [NADH] (EC 1.3.1.9) (PEG.1518) was also identified.

The JB150 genome bears the genes for three major membrane phospholipids phosphatidylethanolamine (PE), phosphatidylglycerol (PG), cardiolipin (CL). To synthesize PE, the strain comprises phosphatidylserine synthase (Pss) (CDP-diacylglycerol--serine O-phosphatidyltransferase) (EC 2.7.8.8) (PEG.5939) and phosphatidylserine decarboxylase (Psd) (EC 4.1.1.65) (PEG.5940). The genetic determinant for CDP-diacylglycerol-glycerol-3-phosphate 3-phosphatidyltransferase (EC 2.7.8.5) (PEG.5321) was also identified in JB150 that catalyzes the necessary steps in the synthesis of the membrane lipids PG and CL. To synthesize PG, the JB150 genome bears L-O-lysylphosphatidylglycerol synthase (EC 2.3.2.3) (PEG.4026) and cardiolipin synthetase (*cls*) (EC 2.7.8.41) (PEG.1131) (Table S1, Supporting Information). In arid desert regions, to resist extreme fluctuations of temperature, nutrient limitation, desiccation, and water supply, the strain JB150 appears to store primary storage lipophilic compounds, polyhydroxyalkanoates (PHA) and triacylglycerols (TAG) [153,154]. TAG accumulation may confer the strain the capability to utilize this reserve to survive and counteract harsh environmental conditions [155, 156]. For TAG metabolism, we detected triacylglycerol lipase precursor (EC 3.1.1.3) (PEG.6700) in JB150. Similarly, poly-β-hydroxybutyric acid (PHB) granules are stored in the cells as a carbon and energy source when environmental conditions are sub-optimal for survival [157,158]. In JB150, to catalyze the degradation of PHB, intracellular poly(3-hydroxyalkanoate) depolymerase (EC 3.1.1.-) (PEG.604), D-β-hydroxybutyrate dehydrogenase (EC 1.1.1.30) (PEG.647) and 3-ketoacyl-CoA thiolase (EC 2.3.1.16) (PEG.5563) were identified (Table S1a, Supporting Information). The comparison among JB150 and the other analyzed genomes reveals the presence of a large number of genes (38) of glycerolipid and glycerophospholipid metabolism in the genome of *L. deserti* while these genes were observed to be absent in *J. gansuensis*. The genes for carotenoids synthesis were shared by JB150 and the other 2 analyzed *Streptomyces* genomes and *A. spitiensis*. All the genomes have been found to possess multiple copies of polyhydroxybutyrate metabolism genes, the highest number of genes (42) of which were observed to be possessed by *L. deserti* (Table S3, Supporting Information).

### Cofactors, vitamins, prosthetic groups, pigments, and motility

The genome of *Streptomyces* JB150 possesses 168 genes (13.5%) for cofactors, vitamins, prosthetic groups, and pigments. The largest gene family includes 47 genes in the subsystem and was observed to be associated with biotin synthesis (Fig.8). Biotin is an essential cofactor for metabolic enzymes such as biotin carboxylase of acetyl-CoA carboxylase (EC 6.3.4.14) (PEG.4524) [164]. The biosynthesis of biotin in JB150 possibly involves the functions of 8-amino-7-oxononanoate synthase (EC 2.3.1.47) (PEG.911) and biotin synthase (EC 2.8.1.6) (PEG.912) which catalyzes the first and final steps in the conserved biosynthetic pathway. The presence of adenosylmethionine-8-amino-7-oxononanoate aminotransferase (EC 2.6.1.62) (PEG.913) (8-amino-7-oxononanoate synthase analog) and dethiobiotin synthase (BioD) (EC 6.3.3.3) (PEG.914) (Table S1a, Supporting Information) suggest further their imperative roles in biotin biosynthetic pathways [159,160]. The negative bifunctional regulatory protein, biotin-protein ligase (EC 6.3.4.9) (PEG.4531), previously identified in the biotin operon in *E. coli* [161], was also observed. Based on the analysis of the co-occurrence of the biotin biosynthetic genes and BioY (PEG.2469) (a substrate-specific component of biotin ECF transporter), it can be predicted that this transmembrane protein is involved in an efficient biotin transport in JB150 (Table S1a, Supporting Information). The other most prominent members identified in the subsystem were the genes involved in the metabolism of riboflavin, flavin mononucleotide (FMN), and flavin adenine dinucleotide (FAD). Riboflavin (vitamin B2) is an ultimate precursor in the biosynthesis of the highly conserved and indispensable flavonoid redox cofactors like FMN and FAD [162,163]. These may serve as cofactors for flavoenzymes in various biochemical reactions, including electron transfer at redox centers, dehydrogenation, oxidation, and hydroxylation reactions [165,166]. The comparison of JB150 and other 7 analyzed genomes reveals that *Streptomyces* spp. JB150, Wb2n-11, and DH-12 possess an unusually high numbers of riboflavin (25), FMN (27) and FAD (27) genes (Table S3, Supporting Information) However, compared to other genomes, the genes for quinone cofactors (menaquinone) were identified only in non-streptomycete genomes which might be contributing their roles in the respiratory electron transport chain [167,168]. Notably, the identification of menaquinone biosynthesis (UbiD family decarboxylase) (PEG.4157), menaquinone polyprenyltransferase (MenA homolog) (PEG.4158), transcriptional regulator (AsnC family) (PEG.941), demethylmenaquinone methyltransferase (EC 2.1.1.163) (PEG.4187) proteins in JB150 indicates the presence of the alternative futalosine pathway [169]. All the analyzed genomes contain the necessary genes to encode essential enzymes involved in folate and pterines synthesis and thus these strains could be predicted as prototrophic for these cofactors. These essential enzymes include GTP cyclohydrolase I (EC 3.5.4.16) (PEG.4020), dihydroneopterin aldolase (EC 4.1.2.25) (PEG.4023), dihydropteroate synthase (EC 2.5.1.15) (PEG.4025), pterin-4-alpha-carbinolamine dehydratase (EC 4.2.1.96) (PEG.6008). Notably, the highest copies of 5-formyltetrahydrofolate cyclo-ligase (EC 6.3.3.2) genes (40) were identified in the genome of *M. excelsi* and *G. africanus* which seem to further support these strains in folate-mediated one-carbon metabolism [170]. Under nutrient-poor conditions, actinomycetes enter into the developmental processes that lead to the formation of spores. The gray-pigmented and rounded spores of *Streptomyces* spp. have thick cell walls and are better adapted to harsh edaphic conditions of the desert [82]. However, compared to other analyzed genomes, the biosynthetic gene cluster which encodes spore pigment was observed to be absent in *J. gansuensis* (Table S3, Supporting Information). The genes for carotenoids synthesis were commonly shared by JB150 and the other two analyzed *Streptomyces* genomes and *A. spitiensis*. Furthermore, the comparison of all the analyzed genomes revealed one interesting feature associated with the genomes of *G. africanus* [11] and *M. excelsi* [14]. These genomes shared 35 genes for flagellar motility. In the genome of *M. excelsi*, a protein lytic transglycosylase, clustered with flagellum synthetic genes (PEG.4945) was detected (Table S3, Supporting Information). The motility in actinomycetes has been detected in the holocarpic genus *Dermatophilus* [171] and in the sporangia-forming *Actinoplanes*, *Ampullariella*, and *Spirillospora* (Couch 1963). The motility elements in these actinomycetes may display a positive chemotactic response to various sugars, inorganic substrates, amino acids, and aromatic compounds [173, 174].

## Conclusion

The urgent need for novel antibiotics to treat multidrug-resistant pathogens has shifted the attention of many researchers to recruit specialized niche like deserts for the identification of actinomycete isolates for their yet unidentified and therapeutically useful secondary metabolites. *Streptomyces* sp. JB150 native to the great Indian Thar desert is capable of growing under such an ecosystem. The whole-genome analysis of JB150 revealed that it encompasses unique gene clusters to produce diverse metabolic compounds. Under extreme conditions, JB150 appears to regulate its genetic machinery to integrate multiple signals to exert appropriate responses to multiple stressors. The presence of many classical as well as distinctive BGCs confers upon the strain the ability to develop several adaptation strategies. The genes required for the synthesis of secondary metabolites were observed to be clustered together at both ends of the JB150 chromosome and exhibit tight regulation. Among these, various BGCs encode for prospective novel chemical scaffolds which can be utilized to target multi-drug resistant microbial pathogens. The genome analysis and metabolic mapping of JB150 revealed prototrophy for many essential amino acids and enrichment of genes involved in various metabolic pathways and protein synthesis. The presence of these genetic attributes in *Streptomyces* sp. JB150 confirms its metabolic versatility to deal with the harsh conditions in the desert environment and oligotrophic nutrient conditions. The comparison of the genomes of JB150 and other taxonomically related desert actinomycetes unveiled the widespread presence of many distinctive and specialized genomic signatures which may assist these strains to encode various antibiotics, antioxidants, CAZymes families, chaperones, defense molecules, DNA repair enzymes, osmoprotectants, proton pumps, siderophores, transporters, etc. The metabolic adaptation and resilience of these strains could provide them a selective advantage over the other microbes. In conclusion, the genetic traits endowed by JB150 and other 7 analyzed genomes could enable them to produce a diverse range of secondary metabolites and stress-responsive proteins to compete, survive and thrive in the desert environment against multiple biotic as well as abiotic stresses.

## Supplementary information

## Additonal file 1

Complementary analyses conducted in the present work on the analyzed genomes.

a. RAST annotations for the *Streptomyces* sp. JB150 genome
b. PATRIC annotations for the *Streptomyces* sp. JB150 genome
c. Predicted secondary metabolites gene clusters from *Streptomyces* sp. JB150 genome
d. Membrane transporter families in *Streptomyces* sp. JB150 genome
e. Clusters of orthologous genes (COGs) of proteins in the genomes of *Streptomyces* sp. JB150 and other 7 desert-actinomycetes, *Streptomyces* sp. Wb2n-11, *Streptomyces* sp. DH-12, *G. africanus*, *L. deserti*, *J. gansuensis*, *M. excelsi* 1G6, and *A. spitiensis*
f. Carbohydrate-active enzymes (CAZymes) families in *Streptomyces* sp. JB150 genome
g. Cross-genus protein families (PGfams) shared by the genomes of *Streptomyces* sp. JB150 and other seven desert-actinomycetes, *Streptomyces* sp. Wb2n-11, *Streptomyces* sp. DH-12, *G. africanus*, *L. deserti*, *J. gansuensis*, *M. excelsi* 1G6, and *A. spitiensis*
h. Unique (orphan) genes in the genome of *Streptomyces* sp. JB150
i. The cross-genera protein families (PGfams) shared by JB150 with *Streptomyces* sp. Wb2n-11, *Streptomyces* sp. DH-12, *G. africanus*, *L. deserti*, *J. gansuensis*, *M. excelsi* 1G6, and *A. spitiensis*
j. Specialty genes in the genomes of *Streptomyces* sp. JB150 and other seven desert-actinomycetes, *Streptomyces* sp. Wb2n-11, *Streptomyces* sp. DH-12, *G. africanus*, *L. deserti*, *J. gansuensis*, *M. excelsi* 1G6, and *A. spitiensis*

## Additonal file 2

Comparison of proteomes between JB150 and the other seven complete genomes (*Streptomyces* sp. Wb2n-11, *Streptomyces* sp. DH-12, *G. africanus*, *L. deserti*, *J. gansuensis*, *M. excelsi* 1G6, and *A. spitiensis*).

## Additonal file 3

Sub-systems features of all the analyzed genomes.

## Compliance with ethical standards Competing interests

The authors declare that they have no conflict of interest

## References

1. Koch AL (2003) Were Gram-positive rods the first bacteria? Trends Microbiol 11:166–70.

2. Battistuzzi FU, Hedges SB (2009) A major clade of prokaryotes with ancient adaptations to life on land. Mol Biol Evol 26:335–43.

3. Bérdy J (2012) Thoughts and facts about antibiotics: where we are now and where we are heading. J Antibiot 65:385–95.

4. Bérdy J (2015) Microorganisms producing antibiotics. In: Antibiotics–Current Innovations and Future Trends. Caister Academic Press, Norfolk 49–64.

5. Barbe V et al (2011) Complete genome sequence of *Streptomyces cattleya* NRRL 8057, a producer of antibiotics and fuorometabolites. J Bacteriol 193:5055–5056.

6. Baranasic D, Gacesa R, Starcevic A, Zucko J, Blažič M, Horvat M, Gjuračić K, Fujs Š, Hranueli D, Kosec G, Cullum J (2013) Draft genome sequence of *Streptomyces rapamycinicus* strain NRRL 5491, the producer of the immunosuppressant rapamycin. Genome Announc 29:1.

7. Thumar JT, Dhulia K, Singh SP (2010) Isolation and partial purification of an antimicrobial agent from halotolerant alkaliphilic *Streptomyces aburaviensis* strain Kut-8. World J Microbiol Biotechnol 26:2081–7.

8. Harwani D (2013) Biodiversity of rare thermophilic actinomycetes in the great Indian Thar desert: an overview. Indo Am J Pharm Res 3:934–9.

9. Begani J, Lakhani J, Harwani D (2018) Current strategies to induce secondary metabolites from microbial biosynthetic cryptic gene clusters. Ann Microbiol 68:419–32.

10. Singla AK, Mayilraj S, Kudo T, Krishnamurthi S, Prasad GS, Vohra RM (2005) *A. spitiensis* sp. nov., a novel actinobacterium isolated from a cold desert of the Indian Himalayas. Int J Syst Evol Microbiol 55:2561–2564.

11. Montero-Calasanz Mdel C, Göker M, Pötter G, Rohde M, Spröer C, Schumann P, Gorbushina AA, Klenk HP (2013) *G. africanus* sp. nov., a halotolerant actinomycete isolated from Saharan desert sand. Antonie Van Leeuwenhoek 104:207–216.

12. Jiao JY, Carro L, Liu L, Gao XY, Zhang XT, Hozzein WN, Lapidus A, Huntemann M, Reddy T, Varghese N, Hadjithomas M, Ivanova NN, Göker M, Pillay M, Eisen JA, Woyke T, Klenk HP, Kyrpides NC, Li WJ (2017) Complete genome sequence of *Jiangella gansuensis* strain YIM 002^T^ (DSM 44835^T^), the type species of the genus *Jiangella* and source of new antibiotic compounds. Stand Genomic Sci J 12:21.

13. Camas M, Veyisoglu A, Tatar D, Saygin H, Cetin D, Sazak A, Guven K and Sahin N (2013) *Lechevalieria nigeriaca* sp. nov., isolated from arid soil. Int J Syst Evol Microbiol 63: 3750–3754.

14. Golinska P, Montero-Calasanz MDC, Świecimska M, et al (2020) *Modestobacter excelsi* sp. nov., a novel actinobacterium isolated from a high altitude Atacama Desert soil. Syst Appl Microbiol 43:126051.

15. Jiao J, Paterson J, Busche T, Rückert C, Kalinowski J, Harwani D, Gross H (2018) Draft genome sequence of Streptomyces sp. strain DH-12, a soilborne isolate from the Thar Desert with broad-spectrum antibacterial activity. Genome Announc 6:e00108–18.

16. Köberl M, White RA, Erschen S, El-Arabi TF, Jansson JK, Berg G (2015) Draft Genome Sequence of *Streptomyces* sp. Strain Wb2n-11, a Desert Isolate with Broad-Spectrum antagonism against Soilborne Phytopathogens. Genome Announc 3:e00860–15.

17. Cybulski JS, Clements J, Prakash M (2014) Foldscope: origami-based paper microscope. PloS one 9:e98781.

18. Li JT, Yang J, Chen DC, Zhang XL, Tang ZS (2007) An optimized mini-preparation method to obtain high-quality genomic DNA from mature leaves of sunflower. Genet Mol Res 6:1064–71.

19. Stackebrandt E, Liesack W, Goebel BM (1993) Bacterial diversity in a soil sample from a subtropical Australian environment as determined by 16S rDNA analysis. FASEB J 7:232–6.

20. Mao J, Wang J, Dai HQ, Zhang ZD, Tang QY, Ren B, Yang N, Goodfellow M, Zhang LX, Liu ZH (2011) *Yuhushiella deserti* gen. nov., sp. nov., a new member of the suborder *Pseudonocardineae*. Int J Sys Evol Microbiol 61:621–30.

21. Claverías FP, Undabarrena AN, González M, Seeger M, Cámara BP (2015) Culturable diversity and antimicrobial activity of actinobacteria from marine sediments in Valparaíso bay, Chile Front Microbiol 6:737.

22. Wright ES, Yilmaz LS, Noguera DR (2012) DECIPHER, a search-based approach to chimera identification for 16S rRNA sequences. Appl Environ Microbiol 78:717–25.

23. Saitou N, Nei M (1987) The neighbor-joining method: A new method for reconstructing phylogenetic trees. Mol Biol Evol 4:406–425.

24. Kumar S, Stecher G, Li M, Knyaz C, Tamura K (2018) MEGA X: Molecular evolutionary genetics analysis across computing platforms. Mol Biol Evol 35:1547–1549.

25. Seemann T (2014) Prokka: rapid prokaryotic genome annotation. Bioinformatics. 30:2068–9.

26. Tatusova T, DiCuccio M, Badretdin A, Chetvernin V, Nawrocki EP, Zaslavsky L, Lomsadze A, Pruitt KD, Borodovsky M, Ostell J (2016) NCBI prokaryotic genome annotation pipeline. Nucleic Acid Res 44:6614–24.

27. Benson DA, Cavanaugh M, Clark K, Karsch-Mizrachi I, Ostell J, Pruitt KD, Sayers EW (2018) GenBank. Nucleic Acid Res 46:D41–7.

28. Brettin T, Davis JJ, Disz T, Edwards RA, Gerdes S, Olsen GJ, Olson R, Overbeek R, Parrello B, Pusch GD, Shukla M (2015) RASTtk: a modular and extensible implementation of the RAST algorithm for building custom annotation pipelines and annotating batches of genomes. Sci Rep 5:8365.

29. Wattam AR, Davis JJ, Assaf R, Boisvert S, Brettin T, Bun C, Conrad N, Dietrich EM, Disz T, Gabbard JL, Gerdes S (2017) Improvements to PATRIC, the all-bacterial bioinformatics database and analysis resource center. Nucleic Acids Res 45:D535–42.

30. Blin K, Shaw S, Steinke K, Villebro R, Ziemert N, Lee SY, Medema MH, Weber T (2019) antiSMASH 5.0: updates to the secondary metabolite genome mining pipeline. Nucleic Acid Res 47:W81–7.

31. Gao F, Zhang C (2008) Ori-Finder: A web-based system for finding oriC s in unannotated bacterial genomes. BMC Bioinformatics 9:79.

32. Xu L, Dong Z, Fang L, Luo Y, Wei Z, Guo H, Zhang G, Gu YQ, Coleman-Derr D, Xia Q, Wang Y (2019) OrthoVenn2: a web server for whole-genome comparison and annotation of orthologous clusters across multiple species. Nucleic Acids Res 47:W52–8.

33. van Heel AJ, de Jong A, Montalban-Lopez M, Kok J, Kuipers OP (2013) BAGEL3: automated identification of genes encoding bacteriocins and (non-) bactericidal posttranslationally modified peptides. Nucleic Acids Res 41:W448–53.

34. Skinnider MA, Dejong CA, Rees PN, Johnston CW, Li H, Webster AL, Wyatt MA, Magarvey NA (2015) Genomes to natural products prediction informatics for secondary metabolomes (PRISM). Nucleic Acids Res 43:9645–62.

35. Henry CS. DeJongh M, Best AA, Frybarger PM, Linsay B, Stevens RL (2010) High-throughput generation, optimization and analysis of genome-scale metabolic models. Nat Biotechnol 28:977–82.

36. Takamatsu S, Lin X, Nara A, Komatsu M, Cane DE, Ikeda H (2011) Characterization of a silent sesquiterpenoid biosynthetic pathway in *Streptomyces avermitilis* controlling epi-isozizaene albaflavenone biosynthesis and isolation of a new oxidized epi-isozizaene metabolite. Microb biotechnol 4:184–91.

37. Song C, Mazzola M, Cheng X, Oetjen J, Alexandrov T, Dorrestein P, Watrous J, van der Voort M, Raaijmakers (2015) Molecular and chemical dialogues in bacteria-protozoa interactions. Sci Rep 5:12837.

38. Yamada Y, Kuzuyama T, Komatsu M, Shin-ya K, Omura S, Cane DE, Ikeda H (2015) Terpene synthases are widely distributed in bacteria. Proc Natl Acad Sci 112:857–62.

39. Chen X, Köllner TG, Jia Q, Norris A, Santhanam B, Rabe P, Dickschat JS, Shaulsky G, Gershenzon J, Chen F (2016) Terpene synthase genes in eukaryotes beyond plants and fungi: Occurrence in social amoebae. Proc Natl Acad Sci 113:12132–7.

40. Allam NG, Abd El-Zaher EH (2012) Protective role of *Aspergillus fumigatus* melanin against ultraviolet (UV) irradiation and *Bjerkandera adusta* melanin as a candidate vaccine against systemic candidiasis. Afr J Biotechnol 11:6566–77.

41. Gadd GM, de Rome L (1988) Biosorption of copper by fungal melanin. Appl Microbiol Biot 29: 610–617.

42. Langfelder K, Streibel M, Jahn B, Haase G, Brakhage AA (2003) Biosynthesis of fungal melanins and their importance for human pathogenic fungi. Fungal Genet Biol 38:143–58.

43. Galinski EA, Pfeiffer HP, Trüper HG (1985) 1,4,5,6-Tetrahydro-2-methyl-4-pyrimidinecarboxylic acid. A novel cyclic amino acid from halophilic phototrophic bacteria of the genus *Ectothiorhodospira*. Eur J Biochem 149:135–139.

44. Graf R, Anzali S, Buenger J, Pfluecker F, Driller H (2008) The multifunctional role of ectoine as a natural cell protectant. Clin Dermatol 26:326–33.

45. Harishchandra RK, Wulff S, Lentzen G, Neuhaus T, Galla HJ (2010) The effect of compatible solute ectoines on the structural organization of lipid monolayer and bilayer membranes. Biophy Chem 150:37–46.

46. Roychoudhury A, Haussinger D, Oesterhelt F (2012) Effect of the compatible solute ectoine on the stability of the membrane proteins. Protein Pept Lett 19:791–4.

47. Sun C, Yang Z, Zhang C, Liu Z, He J, Liu Q, Zhang T, Ju J, Ma J (2019) Genome mining of *Streptomyces atratus* SCSIO ZH16: discovery of atratumycin and identification of its biosynthetic gene cluster. Org Lett 21:1453–7.

48. Saugar I, Sánz E, Rubio MA, Espinosa JC, Jiménez A (2002) Identification of a set of genes involved in the biosynthesis of the aminonucleoside moiety of antibiotic A201A from *Streptomyces capreolus*. Eur J Biochem 269:5527–5535.

49. Miao V, Coeffet-LeGal MF, Brian P, Brost R, Penn J, Whiting A, Martin S, Ford R, Parr I, Bouchard M, Silva CJ (2005) Daptomycin biosynthesis in *Streptomyces roseosporus*: cloning and analysis of the gene cluster and revision of peptide stereochemistry. Microbiol 151:1507–23.

50. Zhang C, Sheng C, Wang W, Hu H, Peng H, Zhang X (2015) Identification of the lomofungin biosynthesis gene cluster and associated flavin-Dependent monooxygenase gene in *Streptomyces lomondensis* S015. PLoS One 10:e0136228.

51. Sherman DH, Malpartida F, Bibb MJ, Kieser HM, Bibb MJ, Hopwood DA (1989) Structure and deduced function of the granaticin-producing polyketide synthase gene cluster of *Streptomyces violaceoruber* Tü22. EMBO J 8:2717–2725.

52. Zhang Y, Huang H, Chen Q, Luo M, Sun A, Song Y, Ma J, Ju J (2013) Identification of the grincamycin gene cluster unveils divergent roles for GcnQ in different hosts, tailoring the L-rhodinose moiety. Org Lett 15:3254–7.

53. Ohnishi Y, Ishikawa J, Hara H, Suzuki H, Ikenoya M, Ikeda H, Yamashita A, Hattori M, Horinouchi S (2008) Genome sequence of the streptomycin-producing microorganism *Streptomyces griseus* IFO 13350. J Bacteriol 190:4050–60.

54. Inahashi Y, Zhou S, Bibb MJ, Song L, Al-Bassam MM, Bibb MJ, Challis GL (2017) Watasemycin biosynthesis in *Streptomyces venezuelae*: thiazoline C-methylation by a type B radical-SAM methylase homologue. Chem Sci J 8:2823–31.

55. Tietz JI, Schwalen CJ, Patel PS, Maxson T, Blair PM, Tai HC, Zakai UI, Mitchell DA (2017) A new genome-mining tool redefines the lasso peptide biosynthetic landscape. Nature chem boil 13:470–8.

56. Matsuda K, Hasebe F, Shiwa Y, Kanesaki Y, Tomita T, Yoshikawa H, Shin-ya K, Kuzuyama T, Nishiyama M (2017) Genome mining of amino group carrier protein-mediated machinery: discovery and biosynthetic characterization of a natural product with unique hydrazone unit. ACS Chem Biol 12:124–31.

57. Schnell N, Entian KD, Schneider U, Götz F, Zahner H, Kellner R, Jung G (1988) Prepeptide sequence of epidermin, a ribosomally synthesized antibiotic with four sulphide-rings. Nature 333: 276–278.

58. Maksimov MO, Pan SJ, James Link A (2012) Lasso peptides: structure, function, biosynthesis, and engineering. Nat Prod Rep 29:996–1006.

59. Arnison PG, Bibb MJ, Bierbaum G, Bowers AA, Bugni TS, Bulaj G, Camarero JA, Campopiano DJ, Challis GL, Clardy J, Cotter PD (2013) Ribosomally synthesized and post-translationally modified peptide natural products: overview and recommendations for a universal nomenclature. Nat Prod Rep 30:108–60.

60. Li Y, Zirah S, Rebuffat S (2015) Bacterial Strategies to Make and Maintain Bioactive Entangled Scaffolds. In: Springer Briefs in Microbiology, Springer-Verlag New York.

61. Hegemann JD, Zimmermann M, Xie X, Marahiel MA (2015) Lasso peptides: an intriguing class of bacterial natural products. Acc Chem Res 48:1909–1919.

62. Cox CL, Doroghazi JR, Mitchell DA(2015) The genomic landscape of ribosomal peptides containing thiazole and oxazole heterocycles. BMC Genomics 16:778.

63. Imlay JA (2008) Cellular defenses against superoxide and hydrogen peroxide. Annu Rev Biochem 77:755–76.

64. Chiang SM, Schellhorn HE (2012) Regulators of oxidative stress response genes in *Escherichia coli* and their functional conservation in bacteria. Arch Biochem Biophys 525:161–9.

65. Hahn JS, Oh SY, Roe JH (2002) Role of OxyR as a peroxide-sensing positive regulator in *Streptomyces coelicolor* A3 (2). J Bacteriol 184:5214–22.

66. Imlay JA (2015) Transcription factors that defend bacteria against reactive oxygen species. Annu Rev Microbiol 69:93–108.

67. Fraser CM, Gocayne JD, White O, Adams MD, Clayton RA, Fleischmann RD, Bult CJ, Kerlavage AR, Sutton G, Kelley JM, Fritchman JL (1995) The minimal gene complement of *Mycoplasma genitalium*. Science 270:397–404.

68. Hahn MY, Bae JB, Park JH, Roe JH (2003) Isolation and characterization of *Streptomyces coelicolor* RNA polymerase, its sigma, and antisigma factors. In: Methods in enzymology. Academic Press 73–82.

69. Lonetto MA, Gribskov M, Gross CA (1992) The sigma 70 family: sequence conservation and evolutionary relationships. J Bacteriol 174:3843.

70. Merrick MJ (1993) In a class of its own-the RNA polymerase sigma factor σ; 54 (σN). Mol Microbiol 10:903–9.

71. Studholme DJ, Dixon R (2003) Domain architectures of σ54-dependent transcriptional activators. J Bacteriol 185:1757–67.

72. Paget MS (2015) Bacterial Sigma Factors and Anti-Sigma Factors: Structure, Function and Distribution. Biomolecules 5:1245–65.

73. Mogensen JE, Otzen DE (2005) Interactions between folding factors and bacterial outer membrane proteins. Mol Microbiol 57:326–46.

74. Saier MH (2000) A functional-phylogenetic classification system for transmembrane solute transporters. Microbiol Mol Biol Rev 64:354–411.

75. Chater KF, Biro S, Lee KJ, Palmer T, Schrempf H (2010) The complex extracellular biology of Streptomyces. FEMS Microbiol Rev 34:171–98.

76. Elbourne LD, Tetu SG, Hassan KA, Paulsen IT (2017) TransportDB 2.0: a database for exploring membrane transporters in sequenced genomes from all domains of life. Nucleic Acid Res 45:D320–4.

77. Higgins C (2001) ABC transporters: physiology, structure and mechanism–an overview. Res Microbiol 152:205–10.

78. Bentley SD, Chater KF, Cerdeño-Tárraga AM, Challis GL, Thomson NR, James KD, Harris DE, Quail MA, Kieser H, Harper D, Bateman A (2002) Complete genome sequence of the model actinomycete *Streptomyces coelicolor* A3 (2). Nature 417:141–7.

79. Jack DL, Paulsen IT, Saier MH (2000) The amino acid/polyamine/organocation (APC) superfamily of transporters specific for amino acids, polyamines and organocations. Microbiology 146:1797–1814.

80. Wong FH, Chen JS, Reddy V, Day JL, Shlykov MA, Wakabayashi ST, Saier Jr MH (2012) The amino acid-polyamine-organocation superfamily. J Mol Microbiol Biotechnol 22:105–13.

81. Berrier C, Coulombe A, Szabo I, Zoratti M, Ghazi A (1992) Gadolinium ion inhibits loss of metabolites induced by osmotic shock and large stretch-activated channels in bacteria. Eur J Biochem 206:559–65.

82. Chater KF, Chandra G (2006) The evolution of development in *Streptomyces* analysed by genome comparisons. FEMS Microbiol Rev 30:651–672.

83. Kleiner D (1985) Bacterial ammonium transport. FEMS Microbiol 32:87–100.

84. Caspari T, Urlinger S (1996) The activity of the gluconate-H+ symporter of *Schizosaccharomyces pombe* cells is down-regulated by d-glucose and exogenous cAMP. FEBS Lett 395:272–6.

85. Govindarajan S, Elisha Y, Nevo-Dinur K, Amster-Choder O (2013) The general phosphotransferase system proteins localize to sites of strong negative curvature in bacterial cells. MBio 4:5.

86. Lyons JA, Parker JL, Solcan N, Brinth A, Li D, Shah ST, Caffrey M, Newstead S (2014) Structural basis for polyspecificity in the POT family of proton-coupled oligopeptide transporters. EMBO Rep 15:886–93.

87. Jack DL, Yang NM, Saier MH (2001) The drug/metabolite transporter superfamily. Eur J Biochem 268:3620–39.

88. Yen MR, Tseng YH, Simic P, Sahm H, Eggeling L, Saier Jr MH (2002) The ubiquitous ThrE family of putative transmembrane amino acid efflux transporters. Res Microbiol 153:19–25.

89. Komorsky-Lovrić Š (1998) A simple method for detection of manganese in marine sediments. Croatica chemica acta 4:263–9.

90. Li W, Tang X (2012) Manganese trafficking, metabolism and homeostasis. Chin Bull Life Sci 24:867–80.

91. Rosen BP. Bacterial resistance to heavy metals and metalloids. J Biol Inorg 1996;1:273–7.

92. Hughes MF (2002) Arsenic toxicity and potential mechanisms of action. Toxicol 133:1–6.

93. Fu HL, Meng Y, Ordóñez E, Villadangos AF, Bhattacharjee H, Gil JA, Mateos LM, Rosen BP (2009) Properties of arsenite efflux permeases (Acr3) from *Alkaliphilus metalliredigens* and *Corynebacterium glutamicum*. J Biol Chem 284:19887–95.

94. Padan E, Schuldiner S (1994) Molecular physiology of the Na+/H+ antiporter in *Escherichia coli*. J Exp Biol 196:443–56.

95. Detmers FJ, Lanfermeijer FC, Abele R, Jack RW, Tampé R, Konings WN, Poolman B (2000) Combinatorial peptide libraries reveal the ligand-binding mechanism of the oligopeptide receptor OppA of *Lactococcus lactis*. Proc Natl Acad Sci 97:12487–92.

96. Doeven MK, Abele R, Tampé R, Poolman B (2004) The binding specificity of OppA determines the selectivity of the oligopeptide ATP-binding cassette transporter. J Biol Chem 279:32301–7.

97. Macomber L, Imlay JA (2009) The iron-sulfur clusters of dehydratases are primary intracellular targets of copper toxicity. Proc Natl Acad Sci 106:8344–9.

98. Rensing C, McDevitt SF (2013) The copper metallome in prokaryotic cells. In: Metallomics and the Cell. Springer, Dordrecht, 417–450.

99. Gaetke LM, Chow-Johnson HS, Chow CK (2014) Copper: toxicological relevance and mechanisms. Arch Toxicol 88:1929–38.

100. Cha JS, Cooksey DA (1991) Copper resistance in *Pseudomonas syringae* mediated by periplasmic and outer membrane proteins. Proc Natl Acad Sci 88:8915–9.

101. Hirooka K, Edahiro T, Kimura K, Fujita Y (2012) Direct and indirect regulation of the *ycnKJI* operon involved in copper uptake through two transcriptional repressors, YcnK and CsoR, in *Bacillus subtilis*. J Bacteriol 194:5675–87.

102. Kobayashi M, Shimizu S (1999) Cobalt proteins. Eur J Biochem 261:1–9.

103. Mulrooney SB, Hausinger RP (2003) Nickel uptake and utilization by microorganisms. FEMS Microbiol Rev 27:239–261.

104. Bogsch EG, Sargent F, Stanley NR, Berks BC, Robinson C, Palmer T (1998) An essential component of a novel bacterial protein export system with homologues in plastids and mitochondria. J Biol Chem 273:18003–6.

105. Sargent F, Bogsch EG, Stanley NR, Wexler M, Robinson C, Berks BC, Palmer T (1998) Overlapping functions of components of a bacterial Sec-independent protein export pathway. EMBO J 17:3640–50.

106. Weiner JH, Bilous PT, Shaw GM, Lubitz SP, Frost L, Thomas GH, Cole JA, Turner RJ (1998) A novel and ubiquitous system for membrane targeting and secretion of cofactor-containing proteins. Cell 93:93–101.

107. Padan E, Bibi E, Ito M, Krulwich TA (2005) Alkaline pH homeostasis in bacteria: new insights. BBA-biomembranes 1717:67–88.

108. Krulwich TA, Sachs G, Padan E (2011) Molecular aspects of bacterial pH sensing and homeostasis. Nat Rev Microbiol 5:330–43.

109. Padan E, Landau M (2016) Sodium-proton (Na+/H+) antiporters: properties and roles in health and disease. In: The Alkali Metal Ions: Their Role for Life. Springer, Cham, 391–458.

110. Preiss L, Hicks DB, Suzuki S, Meier T, Krulwich TA (2015) Alkaliphilic bacteria with impact on industrial applications, concepts of early life forms, and bioenergetics of ATP synthesis. Front Bioeng Biotechnol 3:75.

111. Hamamoto T, Hashimoto M, Hino M, Kitada M, Seto Y, Kudo T, Horikoshi K (1994) Characterization of a gene responsible for the Na+/H+ antiporter system of alkalophilic *Bacillus* species strain C-125. Mol Microbiol 14:939–46.

112. Rodionov DA, Hebbeln P, Eudes A, ter Beek J, Rodionova IA, Erkens GB, Slotboom DJ, Gelfand MS, Osterman AL, Hanson AD, Eitinger T (2009) A novel class of modular transporters for vitamins in prokaryotes. J Bacteriol 191:42–51.

113. Forward JA, Behrendt MC, Wyborn NR, Cross R, Kelly DJ (1997) TRAP transporters: A new family of periplasmic solute transport systems encoded by the *dctPQM* genes of *Rhodobacter capsulatus* and by homologs in diverse Gram-negative bacteria. J Bacteriol 179:5482–5493.

114. Rabus R, Jack D, Kelly DJ, Saier Jr. MH (1999) TRAP Transporters: An ancient family of periplasmic solute-receptor dependent secondary active transporters. Microbiol 145:3431–3445.

115. Fisher SH (1992) Glutamine synthesis in *Streptomyces*-a review. Gene 115:13–7.

116. Tiffert Y, Supra P, Wurm R, Wohlleben W, Wagner R, Reuther J (2008) The *Streptomyces coelicolor* GlnR regulon: identification of new GlnR targets and evidence for a central role of GlnR in nitrogen metabolism in actinomycetes. Mol Microbiol 67:861–80.

117. Tiffert Y, Franz-Wachtel M, Fladerer C, Nordheim A, Reuther J, Wohlleben W, Mast Y (2011) Proteomic analysis of the GlnR-mediated response to nitrogen limitation in *Streptomyces coelicolor* M145. Appl Microbiol Biotechnol 89:1149–59.

118. Humbert R, Simoni RD (1980) Genetic and biomedical studies demonstrating a second gene coding for asparagine synthetase in *Escherichia coli*. J Bacteriol 142:212–20.

119. Reitzer LJ, Magasanik B (1982) Asparagine synthetases of *Klebsiella aerogenes*: properties and regulation of synthesis. J Bacteriol 151:1299–313.

120. Neyrolles O, Ferris S, Behbahani N, Montagnier L, Blanchard A (1996) Organization of *Ureaplasma urealyticum* urease gene cluster and expression in a suppressor strain of *Escherichia coli*. J Bacteriol 178:647–655.

121. Caldara M, Minh PN, Bostoen S, Massant J, Charlier D (2007) ArgR-dependent repression of arginine and histidine transport genes in *Escherichia coli* K-12. J Mol Biol 373:251–267.

122. Taj R, Sorensen JL (2015) Synthesis of actinomycetes natural products JBIR-94, JBIR-125, and related analogues. Tetrahedron Lett 56:7108–11.

123. Krysenko S, Okoniewski N, Kulik A, Matthews A, Grimpo J, Wohlleben W, Bera A (2017) Gamma-glutamylpolyamine synthetase GlnA3 is involved in the first step of polyamine degradation pathway in *Streptomyces coelicolor* M145. Front Microbiol 8:726.

124. Bentley R, Haslam E (1990) The shikimate pathway-a metabolic tree with many branches. Crit Rev Biochem Mol 25:307–84.

125. Cantarel BL, Coutinho PM, Rancurel C, Bernard T, Lombard V, Henrissat B (2009) The Carbohydrate-Active EnZymes database (CAZy): an expert resource for glycogenomics. Nucleic Acid Res 37:D233–8.

126. Henrissat B (1991) A classification of glycosyl hydrolases based on amino acid sequence similarities. Biochem J 280:309–16.

127. Berlemont R, Martiny AC (2016) Glycoside hydrolases across environmental microbial communities. PLoS Comput Biol 12:e1005300.

128. Coutinho PM, Deleury E, Davies GJ, Henrissat B (2003) An evolving hierarchical family classification for glycosyltransferases. J Mol Biol 328:307–17.

129. Lairson LL, Henrissat B, Davies GJ, Withers SG (2008) Glycosyltransferases: structures, functions, and mechanisms. Annu Rev Biochem 77:521–55.

130. Breton C, Šnajdrová L, Jeanneau C, Koča J, Imberty A (2006) Structures and mechanisms of glycosyltransferases. Glycobiology 16:29R–37R.

131. Yip VL, Withers SG (2006) Breakdown of oligosaccharides by the process of elimination. Curr Opin Chem Biol 10:147–55.

132. Christov LP, Prior BA (1993) Esterases of xylan-degrading microorganisms: production, properties, and significance. Enzyme and Microb Technol 15:460–75.

133. Vaaje-Kolstad G, Westereng B, Horn SJ, Liu Z, Zhai H, Sørlie M, Eijsink VG (2010) An oxidative enzyme boosting the enzymatic conversion of recalcitrant polysaccharides. Science 330:219–22.

134. Eijsink VG, Petrovic D, Forsberg Z, Mekasha S, Røhr ÅK, Várnai A, Bissaro B, Vaaje-Kolstad G (2019) On the functional characterization of lytic polysaccharide monooxygenases (LPMOs). Biotechnol Biofuels 12:1–6.

135. Shoseyov O, Shani Z, Levy I (2006) Carbohydrate binding modules: biochemical properties and novel applications. Microbiol Mol Biol Rev 70:283–95.

136. Nishida H, Nishiyama M, Kobashi N, Kosuge T, Hoshino T, Yamane H (1999) A prokaryotic gene cluster involved in synthesis of lysine through the amino adipate pathway: a key to the evolution of amino acid biosynthesis. Genome Res 9:1175–83.

137. Kosuge T, Hoshino T (1998) Lysine is synthesized through the α-aminoadipate pathway in *Thermus thermophilus*. FEMS Microbiol 169:361–7.

138. Kobashi N, Nishiyama M, Tanokura M (1999) Aspartate kinase-independent lysine synthesis in an extremely thermophilic bacterium, *Thermus thermophilus*: lysine is synthesized via α-aminoadipic acid not via diaminopimelic acid. J Bacteriol 181:1713–8.

139. Noens EE, Mersinias V, Traag BA, Smith CP, Koerten HK, van Wezel GP (2005) SsgA-like proteins determine the fate of peptidoglycan during sporulation of *Streptomyces coelicolor*. Mol Microbiol 58:929–44.

140. Liu Y, Prasad R, Beard WA, Kedar PS, Hou EW, Shock DD, Wilson SH (2007) Coordination of steps in single-nucleotide base excision repair mediated by apurinic/apyrimidinic endonuclease 1 and DNA polymerase β. J Biol Chem 282:13532–41.

141. Van der Oost J, Jore MM, Westra ER, Lundgren M, Brouns SJ (2009) CRISPR-based adaptive and heritable immunity in prokaryotes. Trends Biochem Sci 2009;34:401–7.

142. Makarova KS, Haft DH, Barrangou R, Brouns SJ, Charpentier E, Horvath P, Moineau S, Mojica FJ, Wolf YI, Yakunin AF, Van Der Oost J (2011) Evolution and classification of the CRISPR–Cas systems. Nat Rev Microbiol 9:467–77.

143. Barrangou R, Horvath P (2012) CRISPR: new horizons in phage resistance and strain identification. Annu Rev Food Sci T 3:143–62.

144. Wiedenheft B, Sternberg SH, Doudna JA (2012) RNA-guided genetic silencing systems in bacteria and archaea. Nature 482:331–8.

145. Jansen R, Embden JD, Gaastra W, Schouls LM (2002) Identification of genes that are associated with DNA repeats in prokaryotes. Mol Microbiol 43:1565–75.

146. Makarova KS, Koonin EV (2015) Annotation and Classification of CRISPR-Cas Systems. Methods Mol Biol 13:47–75.

147. Dryden DT, Murray NE, and Rao DN (2001) Nucleoside triphosphate-dependent restriction enzymes. Nucleic Acids Res 29:3728–3741.

148. Murray NE (2002) Immigration control of DNA in bacteria: self versus non-self. Microbiol 148: 3–20.

149. Revill WP, Bibb MJ, Hopwood DA (1995) Purification of a malonyltransferase from *Streptomyces coelicolor* A3 (2) and analysis of its genetic determinant. J Bacteriol 177:3946–52.

150. Revill WP, Bibb MJ, Scheu AK, Kieser HJ, Hopwood DA (2001) β-Ketoacyl acyl carrier protein synthase III (FabH) is essential for fatty acid biosynthesis in *Streptomyces coelicolor* A3 (2). J Bacteriol 183:3526–30.

151. Florova G, Kazanina G, Reynolds KA (2002) Enzymes involved in fatty acid and polyketide biosynthesis in *Streptomyces glaucescens*: role of FabH and FabD and their acyl carrier protein specificity. Biochemistry 41:10462–71.

152. Han L, Lobo S, Reynolds KA (1998) Characterization of β-ketoacyl-acyl carrier protein synthase III from *Streptomyces glaucescens* and its role in initiation of fatty acid biosynthesis. J Bacteriol 180:4481–6.

153. Steinbüchel A (2001) Perspectives for biotechnological production and utilization of biopolymers: metabolic engineering of polyhydroxyalkanoate biosynthesis pathways as a successful example. Macromol Biosci 1:1–24.

154. Alvarez H, Steinbüchel A (2002) Triacylglycerols in prokaryotic microorganisms. Appl Microbiol biotechnol 60:367–76.

155. Alvarez HM, Souto MF, Viale A, Pucci OH (2001) Biosynthesis of fatty acids and triacylglycerols by 2, 6, 10, 14-tetramethyl pentadecane-grown cells of *Nocardia globerula* 432. FEMS microbiol lett 200:195–200.

156. Olukoshi ER, Packter NM (1994) Importance of stored triacylglycerols in *Streptomyces*: possible carbon source for antibiotics. Microbiol 140:931–43.

157. Mergaert J, Webb A, Anderson C, Wouters A, Swings J (1993) Microbial degradation of poly (3-hydroxybutyrate) and poly (3-hydroxybutyrate-co-3-hydroxyvalerate) in soils. Appl Environ Microbiol 59:3233–8.

158. Jendrossek D (2009) Polyhydroxyalkanoate granules are complex subcellular organelles (carbonosomes). J Bacteriol 191:3195–202.

159. Lin S, Cronan JE (2011) Closing in on complete pathways of biotin biosynthesis. Mol Biosyst 7:1811–1821.

160. Schneider G, Lindqvist Y (2001) Structural enzymology of biotin biosynthesis. FEBS Lett 495:7–11.

161. DeMoll E (1994) Biosynthesis of biotin and lipoic acid in *Escherichia coli* and *Salmonella*. In: Cellular and molecular biology. American Society for Microbiology, Washington, DC, 704–709.

162. Bacher A, Eberhardt S, Eisenreich W, Fischer M, Herz S, Illarionov B, Kis K, Richter G (2001) Biosynthesis of riboflavin. Vitam Horm 61:1–49.

163. Fischer M, Bacher A (2005) Biosynthesis of flavocoenzymes. Nat Prod Rep 22:324–50.

164. Perkins JB, Pero JG (2001) Bacillus subtilis and Its Relatives. In: From Genes to Cells. American Society for Microbiology, Washington, DC, 279–293.

165. Massey V (1995) Introduction: flavoprotein structure and mechanism. FASEB J 9:473–475.

166. Fraaije W. Mattevi A (2000) Flavoenzymes: diverse catalysts with recurrent features. Trends Biochem Sci 25:126–132.

167. Nitschke W, Kramer DM, Riedel A, Liebl U (1995) From naphtho-to benzoquinones-(r) evolutionary reorganizations of electron transfer chains. In: Photosynthesis: from light to biosphere. Springer, 945–50.

168. Soballe R, Poolle RK (1999) Microbial ubiquinones: multiple roles in respiration, gene regulation and oxidative stress management. Microbiology 145:1817–30.

169. Hiratsuka T, Furihata K, Ishikawa J, Yamashita H, Itoh N, Seto H, Dairi T (2008) An alternative menaquinone biosynthetic pathway operating in microorganisms. Science 32:1670–3.

170. Lan X, Field MS, Stover PJ (2018) Cell cycle regulation of folate-mediated one-carbon metabolism. Wiley Interdiscip Rev Syst Biol Med 10:e1426.

171. Gordon MA (1964) The genus *Dermatophilus*. J Bacteriol 88:509–22.

172. Couch JN (1963) Some new genera and species of the *Actinoplanaceae*. J Elisha Mitchell Sci Soc 79:53–70.

173. Arora DK (1986) Chemotaxis of *Actinoplanes missouriensis* zoospores to fungal conidia, chlamydospores and sclerotia. J Gen Microbiol 132:1657–1663.

174. Hayakawa M, Tamura T, Nonomura H (1991) Selective isolation of *Actinoplanes* and *Dactylosporangium* from soil by using y-collidine as the chemoattractant. J Ferment Bioeng 72:426–432.

